# Multiple *cis*-regulatory elements control *prox1a* expression in distinct lymphatic vascular beds

**DOI:** 10.1101/2023.08.21.550483

**Authors:** Virginia Panara, Hujun Yu, Di Peng, Karin Staxäng, Monika Hodik, Beata Filipek-Gorniok, Jan Kazenwadel, Renae Skoczylas, Elizabeth Mason, Amin Allalou, Natasha L. Harvey, Tatjana Haitina, Benjamin M. Hogan, Katarzyna Koltowska

## Abstract

Lymphatic vessels play a role in several physiological and pathological processes including tissue fluid homeostasis, dietary fat absorption, immunosurveillance, and immunomodulation. During embryonic development, lymphatic endothelial cell (LEC) precursors are distinguished from blood endothelial cells by the expression of the transcription factor Prospero-related homeobox 1 (PROX1). PROX1 is essential for lymphatic vascular network formation in mouse and zebrafish. The initiation of PROX1 expression precedes LEC sprouting and migration, serving as the definitive marker of specified LECs. Despite its crucial role in lymphatic development, the upstream regulation of *PROX1* in LECs remains to be uncovered. SOX18 and COUP-TFII are thought to regulate *Prox1* expression in mice by binding to its promoter region. However, how the specificity of *Prox1* expression to LECs is achieved remains to be studied in detail.

In this study, we analysed evolutionary conservation and chromatin accessibility to identify enhancer sequences located in the proximity of zebrafish *prox1a* active in developing LECs. We confirmed the functional role of the identified sequences through CRISPR/Cas9 mutagenesis of a lymphatic valve enhancer. The deletion of this genomic region results in impaired valve morphology and function. Overall, our results reveal the intricate control of *prox1a* expression through a collection of enhancers. Ray-finned fish-specific distal enhancers drive pan-lymphatic expression, while vertebrate-conserved proximal enhancers refine expression in functionally distinct subsets of lymphatic vessels.

**Graphical Abstract:** 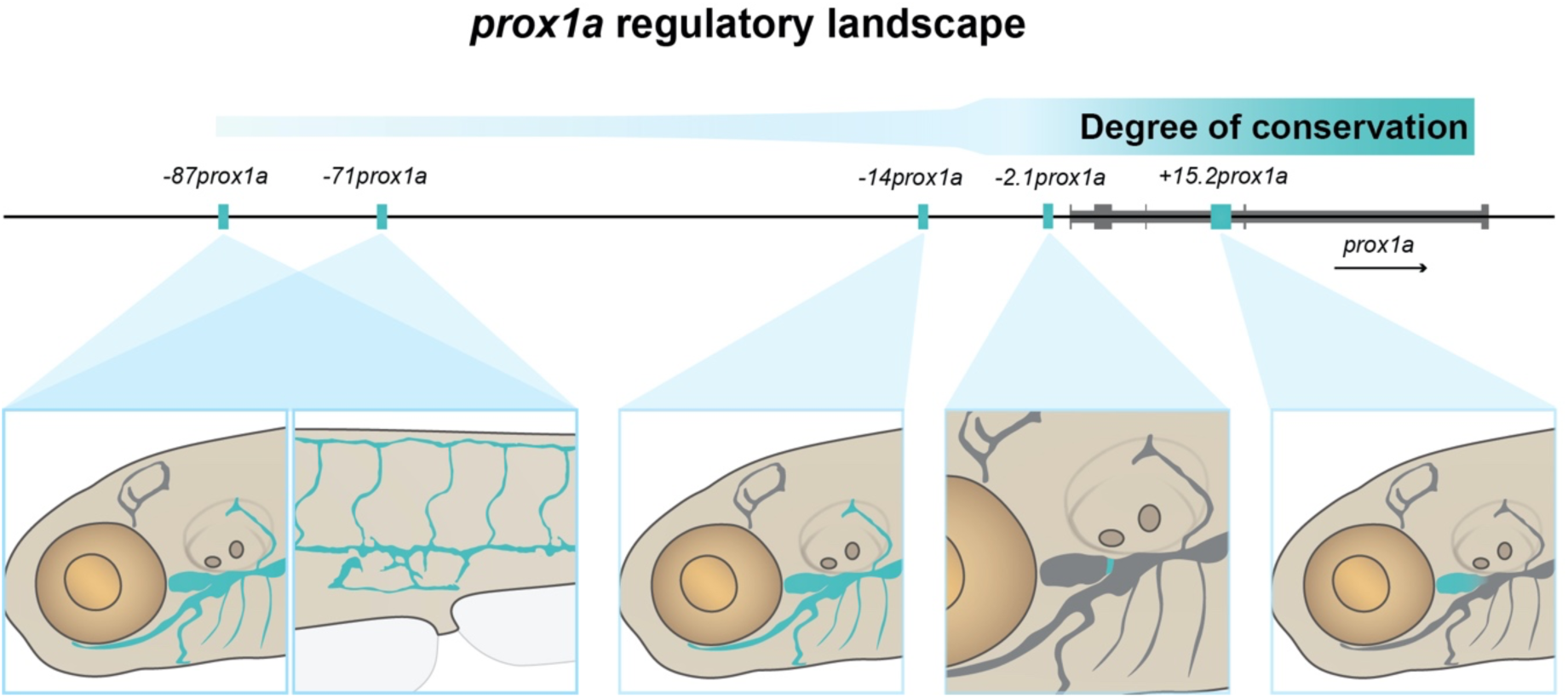

## Introduction

Transcription factor (TF) regulation plays a pivotal role in cell specification and the acquisition of tissue identity during development (Spitz and Furlong, 2012). The TF *PROX1* is a key gene involved in lymphatic vasculature development. In mouse, PROX1 is expressed by the lymphatic progenitors and is the first marker of specified lymphatic endothelial cells (LECs) (Wigle and Oliver, 1999). *Prox1* mutants are characterised by the loss of all lymphatic structures (Wigle and Oliver, 1999). However, during embryonic development, *PROX1* is expressed in multiple tissues, such as the central nervous system, liver, retina and skeletal muscles (Glasgow and Tomarev, 1998; Pistocchi et al., 2008), implying the necessity for mechanisms restricting Prox1 expression in a tissue specific manner. Although TFs regulating *PROX1* expression in the lymphatics have been described in mammals (François et al., 2008; Srinivasan et al., 2010), the upstream *cis-*regulation of *PROX1* remains largely unexplored, leaving a comprehensive understanding of the regulatory logic incomplete.

Enhancers are *cis-*regulatory elements that can be located proximally or distally to a gene locus. Enhancers can function in a modular fashion, with tissue-specific enhancers activated independently (Long et al., 2016). Enhancer regulation is particularly interesting in a developmental context, as many “developmental toolbox” genes are differentially regulated through enhancer activity. Important enhancer elements active in lymphatic and blood vascular development have been identified in genes such as *gata2a* (Shin et al., 2019), *flt1* (Bussmann et al., 2010), *notch1b* (Chiang et al., 2017), *fli1* (Villefranc et al., 2007) and *etsrp* (Veldman and Lin, 2012). However, to date, only one lymphatic-specific *PROX1* enhancer (Kazenwadel et al., 2023), with activity enriched in the lymphatic valves, has been described. Therefore, the number, contextual nature and identity of the enhancers driving *PROX1* expression in lymphatics remains to be investigated.

In zebrafish, the most studied lymphatic beds are the trunk and facial lymphatics (Eng et al., 2019; Hogan et al., 2009; Küchler et al., 2006; Okuda et al., 2012; Yaniv et al., 2006) which require *prox1* for correct development (Grimm et al., 2023). As a result of the teleost-specific genome duplication event, zebrafish have two co-orthologs of mammalian *PROX1*, called *prox1a* and *prox3* (previously referred to as *prox1b*). *prox1a* is an early marker of lymphatic identity, being expressed by lymphatic progenitors in the posterior cardinal vein (PCV) at 32 hours post-fertilization (hpf), prior to the onset of sprouting (Dunworth et al., 2014; Koltowska et al., 2015a). Its expression marks all described lymphatic beds for the duration of larval development. Mutants for *prox1a* and *prox3* show a reduction in the number of lymphatic endothelial cells (LECs) (Grimm et al., 2023), confirming the importance of these factors for LEC development in zebrafish. Importantly, it is *prox1a* that has a prominent developmental role in zebrafish lymphangiogenesis (Grimm et al., 2023; Koltowska et al., 2015a). Despite this, little is known about *prox1a* regulation. Although the TFs COUP-TFII and SOX18 do not seem to regulate *prox1a* (van Impel et al., 2014), a role of VEGFC signalling upstream of *prox1a* in the specification of LECs has been reported (Koltowska et al., 2015a). However, the regulatory elements and enhancer evolution of *prox1a* regulation are still to be described.

In this study, we aimed to characterise the enhancers regulating the expression of *prox1a* in the developing lymphatics. We identified several elements driving expression in different subsets of the lymphatic endothelium. These include both conserved elements across vertebrates and elements specific to actinopterigians. Our results suggest that *prox1a* is tightly regulated in a spatially patterned manner by a cohort of enhancers acting in concert.

## Results

### The prox1a locus is enriched in evolutionary conserved non-coding regions

To identify enhancer elements in the zebrafish *prox1a* locus, we analysed DNA conservation, as Conserved Non-coding Elements (CNE) close to a gene can indicate the presence of enhancers. We aligned the region of the *PROX1/prox1a* locus in eight Osteichthyes species using mVISTA (**Fig. 1A, Table S1**). The species were selected to have a balanced spread along the phylogeny. Local microsynteny was verified by comparing the identities of neighbouring loci (**Fig. S1A**). In all species analysed, a *SMYD2* homolog is located downstream of *prox1a*, while *RPS6KC1* is located upstream of it in all cases except in zebrafish. Consequently, we defined the regions of interest as the sequences between the two loci adjacent to *PROX1/prox1a,* encompassing the long intergenic region upstream of prox1a (300kbp) for all species in the conservation analysis. We recovered 25 conserved non-coding sequences, located both upstream, downstream and in the intronic regions of *PROX1/prox1a* (**Fig. 1A**). We also investigated the conservation in the other *PROX1* co-ortholog, *prox3*. Interestingly, *prox3* presents very low levels of conservation outside of exon sequences, with only two Telost-conserved elements identified in the analysis and none conserved with other vertebrates in intergenic regions (**Fig. S1B**).

**Figure 1:**
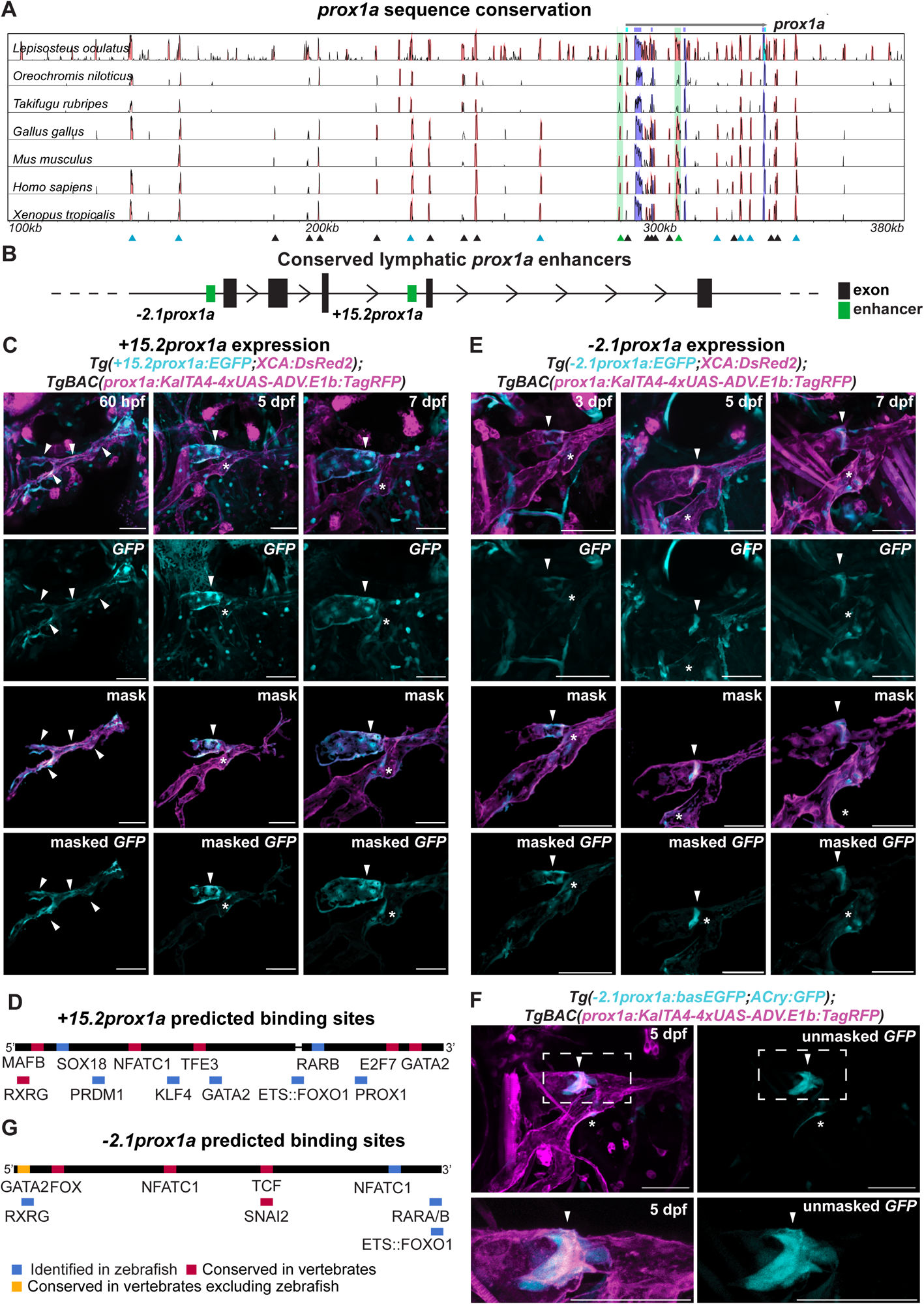
Two conserved *prox1a* enhancers drive expression in subsets of the facial lymphatics. (A) Conservation analysis of the 380 kbp zebrafish region surrounding the *prox1a* locus compared to seven vertebrate species. The results are shown compared to zebrafish. Blue peaks: Exons; red peaks: Region of conserved non-coding DNA; black arrowheads: Conserved peaks; blue arrowheads: Tested peaks; green arrowheads: Identified -2.*1prox1a* and +15.*2prox1a* lymphatic enhancers. In the 5’ the first 100kbp of the alignment contain no conservation peak and has been omitted from the graph. (B) Schematic representation of the *prox1a* locus showing the relative position of the two identified lymphatic enhancers to the *prox1a* exons. Green boxes: the -2.*1prox1a* and +15.*2prox1a* enhancers identified on the basis of sequence conservation; Black boxes: exons. (C) Confocal projections of facial lymphatics labelled with *Tg(+15.2prox1a:EGFP; XCA:DsRed2)^uu7kk^* (cyan) and *Tg(prox1a:RFP)^nim5^* (magenta) at 60 hpf, 5 dpf and 7 dpf. Arrows: expression in the facial LECs (60 hpf) and FCLV (5 and 7 dpf). Asterisks: Expression in facial lymphatic endothelium. (D) Predicted endothelial TF binding sites in *+15.2prox1a*. Blue: Binding sites identified in zebrafish (p-val <1e-04). Red: Conserved binding sites within vertebrates. (E) Confocal projections of facial lymphatics labelled with *Tg(-2.1prox1a:EGFP; XCA:DsRed2)^uu3kk^* (cyan) and *Tg(prox1a:RFP)^nim5^* (magenta) at 3, 5 and 7 dpf. Arrows: Expression in the developing lymphatic valve. Asterisks: Expression in the facial lymphatic endothelium. (F) Confocal projections of the facial lymphatics labelled with *Tg(-2.1prox1a:basEGFP;ACry:GFP)^uu10kk^* (cyan) and *Tg(prox1a:RFP)^nim5^* (magenta) at 5 dpf, top images: Overview of the facial lymphatics, bottom images: zoom of the valve area. Arrow: Expression in the developing lymphatic valve. Asterisk: Expression in the facial lymphatic endothelium. (G) Predicted endothelial TF binding sites in *-2.1prox1a*. Blue: Binding sites identified in zebrafish (p-val < 1e-04). Red: Conserved binding sites within vertebrates. Yellow: Binding sites conserved in vertebrates but absent in zebrafish. In all the images the scale bars represent 50μm.

We then tested selected *prox1a* CNEs for regulatory activity. Histone modifications have been linked to non-coding elements such as promoters and enhancers. Specifically, H3K4me1 marks primed and active enhancers (Heintzman et al., 2007), while H3K27ac indicates active enhancers (Bonn et al., 2012; Creyghton et al., 2010). Using zebrafish public databases for H3K4me1 and H3K27ac, we identified that ten of the selected *prox1a* CNEs were primed or active enhancers (Aday et al., 2011; Bogdanovic et al., 2012) (**Fig. 1A, S1C**). These were subsequently cloned into the Zebrafish Enhancer Detection (ZED) vector (Bessa et al., 2009) and tested *in vivo* in F1 (**Table S2**). Two of the ten tested sequences drove GFP expression in the subsets of lymphatic endothelium. The *+15.2prox1a* element (located 15.2 kbp downstream of the transcription start site) was enriched in the facial collecting lymphatic vessel (FCLV), and -2.*1prox1a* (located 2.1 kbp upstream of the transcription start site) in the lymphatic valve **(Fig. 1B,C,E,F, Table 1, Table S2**).

**Table 1.** *prox1a* enhancers drive regionalised GFP expression in lymphatic endothelium. Quantification of enhancer driven GFP expression across the embryo. The prevalence of expression is marked by +++ highest; ++ medium; + low; - no expression.

| enhancer element | vector type | element length (bp) | Experssion |  |  |  |  |  |  |  | Lymphatic positive/reporter positive identified in F1 |
| --- | --- | --- | --- | --- | --- | --- | --- | --- | --- | --- | --- |
|  |  |  | pan-LEC | Facial LV enriched | Trunk LV enriched | Valve enriched | FCLV enriched | venous | arterial | non-endothelial |  |
| +15.2 prox1 | ZED vector | 927 | - | + | - | + | +++ | ++ | - | ++ | 2/2 |
| -2.1 prox1 | ZED vector | 282 | - | + | - | +++ | - | - | - | ++ | 7/7 |
| -2.1 prox1 | basGFP | 282 | - | + | - | +++ | - | - | - | - | 12/12 |
| -2.1 prox1_1 | ZED vector | 201 | - | + | - | +++ | - | - | - | + | 4/6 |
| -2.1 prox1_2 | ZED vector | 81 | - | + | - | + | + | - | - | ++ | 2/4 |
| -2.1 prox1_3 | ZED vector | 501 | - | ++ | - | +++ | - | - | - | ++ | 3/3 |
| -14.2 prox1 | ZED vector | 712 | - | ++ | - | + | + | - | - | ++ | 6/6 |
| -71.2 prox1 | ZED vector | 584 | + | + | + | + | + | + | - | ++ | 2/2 |
| -87.2 prox1 | ZED vector | 670 | + | + | + | + | + | - | - | ++ | 5/5 |
| empty | ZED vector | 0 | - | - | - | - | - | + | + | ++ | 10/10 |
| empty | basGFP | 0 | - | - | - | - | - | - | - | - | 4/4 |

### +15.2prox1a drives expression in the FCLV

*+15.2prox1a* is a 927 bp element located in intron 3 of *prox1a* **(Fig. 1B)**. The element is composed of two separate conservation peaks divided by 17 bp **(Fig. 1D)**. At 5 and 7 dpf the element is able to drive GFP expression in the FCLV, with faint additional expression in the facial lymphatics **(Fig. 1C**, **Table 1)**. We also observed the reporter expression at 60 hpf, in the developing facial lymphatic, but not in the ventral aorta lymphangioblast (VA-L) **(Fig. 1C-S4A)**. +15.*2prox1a* is in addition active in the venous endothelium in the primary head sinus **(Fig. S4F)** and in the PCV **(Fig. S1D)**. As this element is highly conserved across the vertebrates, we used to p-value cut-off of 1e-02 for our MEME Suite (Bailey and Elkan, 1994) and TOMTOM (Gupta et al., 2007) to determine the motifs and putative transcription factor binding sides. We identified conserved binding sites for multiple known lymphatic regulators, such as GATA2, NFATC1, MAFB, SOX18, PROX1 (Arnold et al., 2022; Geng et al., 2016; Kazenwadel et al., 2015; Koltowska et al., 2015b; Shin et al., 2019) and the retinoic acid receptors RARB and RXRG (Bowles et al., 2014; Marino et al., 2011). We also found putative sites for transcription factors which have been implied in regulation of *Prox1* expression or are related to endothelial cell identity such as TFE3, ETS::FOXO1, PRDM1 and KLF4 (Dieterich et al., 2015; Niimi et al., 2019; Park et al., 2014; Pham et al., 2007; Tai-Nagara et al., 2020; De Val et al., 2008; Yoshimatsu et al., 2011) (**Fig. 1D, S1G-I, Table S4**). This supports that *+15.2prox1a* is an important driver of *prox1a* expression contributing to the regulatory logic necessary for its correct spatial expression in developing lymphatics.

### -2.1prox1a drives expression in the lymphatic valve across developmental stages

*-2.1prox1a* is a conserved sequence of 282 bp located upstream of the TSS (**Fig. 1B**). We recently reported (Kazenwadel et al., 2023) that this element is active in the developing tissue of the lymphatic valve at 5 dpf, and weakly in the rest of the facial lymphatics (**Fig. 1E**, **Table 1**). However, the functional characterisation of this enhancer and the timing of its expression in zebrafish are yet to be determined. Here we uncovered that at 3 dpf the *-2.1prox1a* element is first active in the site of the future valve formation, preceding the *gata2a* onset of expression (Quillien et al., 2017), and its activity is ongoing until valve maturation at 7 dpf (**Fig. 1E**). As the empty ZED vector induces low level of GFP expression in various tissues (**Table 1, Fig. S1E**) and it uses a *gata2a* minimal promoter (Bessa et al., 2009), we wanted to confirm the specificity of *-2.1prox1a* driven expression. We re-cloned *-2.1prox1a* in a different plasmid, using an *e1b TATA* minimal promoter (Villefranc et al., 2007) which does not drive GFP expression without enhancer sequences (**Fig. S1F, Table 1**). We obtained the same enriched expression in the lymphatic valve and sparse expression in facial lymphatics (**Fig. 1F**, **Table 1**), confirming the predominant activity of this enhancer in the valve.

*-2.1prox1a* presents predicted binding sites for a variety of known lymphatic factors. As previously reported, at the 5’ -2.*1prox1a* contains binding sites for NFATC1 and FOX, previously reported to be conserved between zebrafish and mammals, as well as a GATA2 site conserved in other vertebrates, but not in zebrafish (Kazenwadel et al., 2023) **(Fig. 1G, Table S4).** The GATA2 binding to the mice enhancer has been confirmed with ChIP-seq in lymphatic endothelial cells and is functionally necessary for lymphatic vessel formation in mice (Kazenwadel et al., 2023). The enhancer also includes binding sites for genes involved in vascular development such as forkhead box transcription factors (Niimi et al., 2017; Scallan et al., 2021), TCFs (Cha et al., 2016; Nicenboim et al., 2015), SNAI2 (Hultgren et al., 2020), retinoic acid receptors (Bowles et al., 2014; Marino et al., 2011), and ETS factors (Pham et al., 2007; De Val et al., 2008; Yoshimatsu et al., 2011) (**Fig. 1G, S1H-J, Table S4-S5**), thus further implicating the relevance of this enhancer activity in the lymphatic endothelium.

### Accessible regions of open chromatin at the prox1a locus drive expression in the lymphatics

Using the evolutionary conservation analysis, we have identified enhancers that are active in a subset of lymphatic endothelium, but not pan-lymphatic drivers. As non-conserved enhancers also exist to complement our conservation analysis, we utilised the chromatin accessibility approach, which can indicate the presence of an active enhancer in a tissue. To identify additional *prox1a* lymphatic enhancers we used a previously published single nuclei ATAC-seq (snATAC-seq) database of zebrafish ECs (N=3155) at 4 dpf, which included arterial ECs, venous ECs and LECs (Grimm et al., 2023). From control/wild-type sorted EC, we selected all ECs and endocardium, and then subsetted and re-clustered the data (**Fig. S2A**). Cluster identity was determined using the gene accessibility score of marker genes of lymphatic and blood endothelium (**Fig. S2A**). GO terms analysis revealed an enrichment for terms connected with lymphangiogenesis in the LEC-accessible regions, and enrichment for angiogenesis terms in the less accessible regions (**Fig. S2B**).

We focused on the DNA region surrounding the *prox1a* locus and identified six regions of open chromatin in LECs (**Fig. 2A**). We established stable transgenic lines for three of them: - *87prox1a*, -*71prox1a* and *-14prox1a* (**Table S2**). All drove expression in the lymphatic endothelium (**Table 1**). Noticeably, -*2.1prox1a* was also retrieved by this analysis. We cloned the corresponding chromatin accessibility peak, named -*2.1prox1a_3*, which contained an additional 219 base pairs (bp) compared to *-2.1prox1a.* -*2.1prox1a_3* was able to induce expression in the lymphatic valve and residual expression in the rest of the facial lymphatic vessel (**Fig. S2C-D**). This broader expression in the facial LECs is probably due to additional putative TF binding sites present in the sequence which regulate zebrafish lymphangiogenesis, such as Mafb (Arnold et al., 2022; Koltowska et al., 2015b).

**Figure 2:**
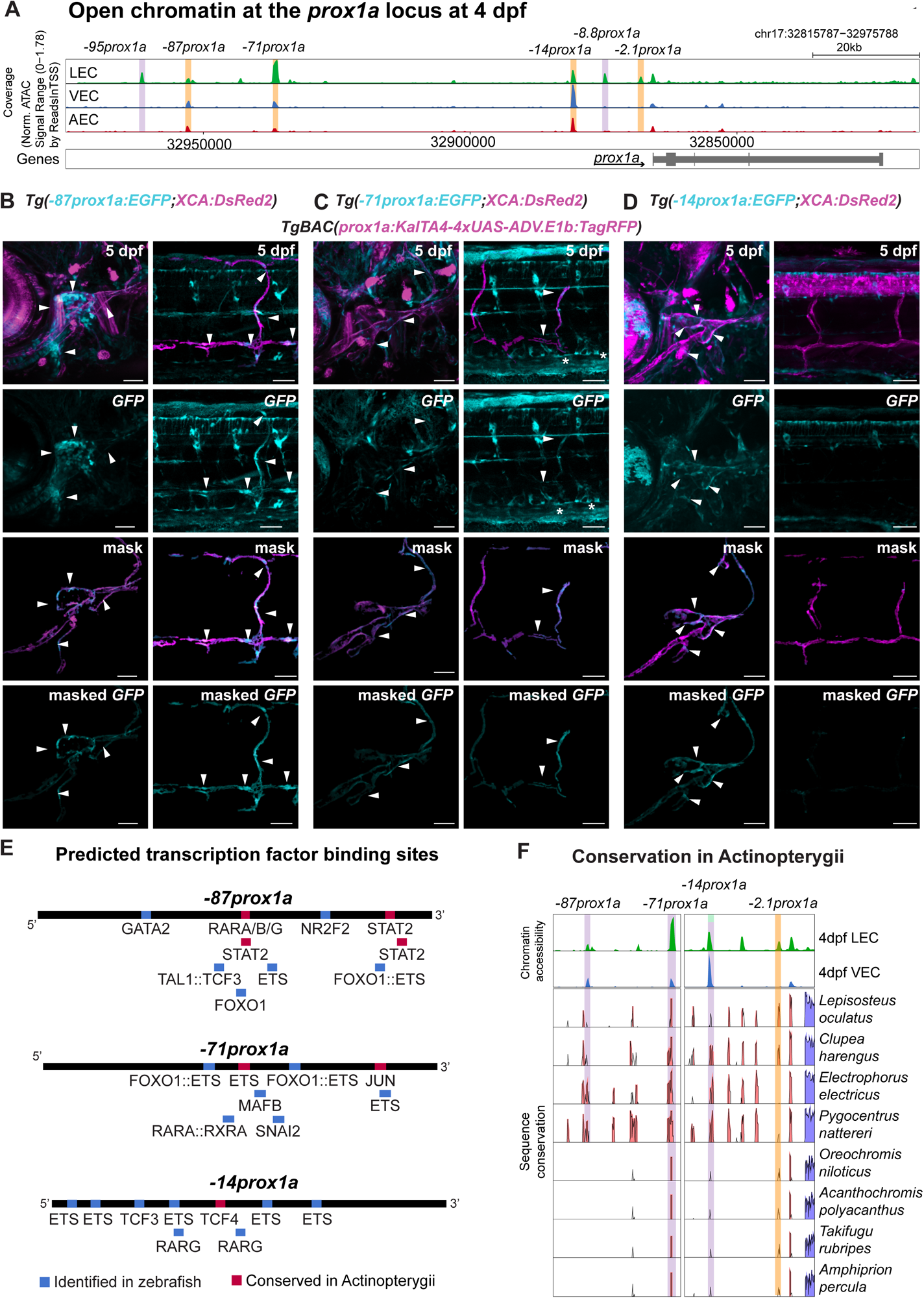
snATAC-seq identifies four lymphatic *prox1a* enhancers. (A) Chromatin state surrounding the *prox1a* locus in lymphatic endothelial cells (LECs) venous endothelial cells (VECs), and arterial endothelial cells (AECs) at 4 dpf, showing the region between 32815787–32975788 base pairs of chromosome (chr) 17. Orange: Tested enhancers. Purple: Identified accessible chromatin sequences in LECs. (B) Confocal projections of the facial and trunk lymphatics labelled with *Tg(-87prox1a:EGFP; XCA:DsRed2)^uom122^* (cyan) and *Tg(prox1a:RFP)^nim5^* (magenta) at 5 dpf. Arrows: Expression in the face and trunk lymphatics. (C) Confocal projections of the facial and trunk lymphatics labelled with *Tg(-71prox1a:EGFP; XCA:DsRed2)^uom121^* (cyan) and *Tg(prox1a:RFP)^nim5^* (magenta) at 5 dpf. Arrows: Expression in the face and trunk lymphatics. Asterisk: Expression in PCV. (D) Confocal projections of the facial and trunk lymphatics labelled with *Tg(-14prox1a:EGFP; XCA:DsRed2)^uom120^* (cyan) and *Tg(prox1a:RFP)^nim5^* (magenta) at 5 dpf. Arrows: Expression in the facial lymphatics. (E) Top: Predicted endothelial TF binding sites in *-87prox1a*. (p-val < 1e-04). Middle: Predicted endothelial TF binding sites in *-71prox1a* (p-val < 1e-04). Bottom: Predicted endothelial TF binding sites in *-14prox1a* (p-val < 1e-04). Blue: Binding sites identified in zebrafish. Red: Conserved binding sites within Actinopterygii. (F) Conservation analysis of the *prox1a* enhancers accessible at 4 dpf in 9 Actinopterygii species. Results are shown compared to zebrafish. Blue peaks: Exons; red peaks: Region of conserved non-coding DNA; Purple bars: Tested enhancer position; Orange bar: *-2.1prox1a* enhancer position.

We further investigated if the enhancers identified by snATAC-seq present sequence conservation within Actinopterygii. Microsynteny was tested in nine ray-finned fish species (**Fig. S3A**) confirming that the loss of *rps6kc1* upstream of *prox1a* we observed in zebrafish is a Otocephala-specific rearrangement and is not present in other Actinopterygii (**Fig. S1A, S3A**). The sequence conservation analysis revealed that *-71prox1a* and *-14prox1a* are conserved across Actinopterygii, while *-87prox1a* could not be identified with sequence conservation in any of the considered acanthopterygian species (*Oreochromis niloticus*, *Acanthochromis polycanthus*, *Takifugu rubipes* and *Amphiprion percula*) (**Fig. 2F).** However, the presence of these enhancers in *Lepisosteus oculatus* suggests the enhancer was present in the ancestor of Actinopterygii, and the sequence has subsequently diverged or was lost in the acanthopterygian lineage.

The *-87prox1a* element, positioned 87 kbp upstream of the TSS, drove reporter expression in the facial and trunk lymphatics at 5 dpf (**Fig. 2B**). Interestingly, it contained predicted TF binding sites for vascular regulators such as GATA2, FOX, ETS, NR2F2, TAL1::TCF3 (Cha et al., 2016; Frye et al., 2018; Kazenwadel et al., 2015; Lin et al., 2010; Nicenboim et al., 2015; Niimi et al., 2017; Pham et al., 2007; Scallan et al., 2021; Tang et al., 2006; De Val et al., 2008; Yoshimatsu et al., 2011), as well as actinopterygians-conserved binding sites for RARA (Bowles et al., 2014; Marino et al., 2011) (**Fig. 2E, S3B, Table S4-S5**).

The *-71prox1a* element is situated 71kbp upstream of the *prox1a* TSS and drove reporter expression in the trunk and the facial lymphatics at 5 dpf (**Fig. 2C**). It contained predicted binding sites for factors involved in vascular development such as MAFB and ETS factors (Arnold et al., 2022; Koltowska et al., 2015b; Pham et al., 2007; De Val et al., 2008; Yoshimatsu et al., 2011) (**Figure 2E, S3C, Table S4-S5**). The enhancer also presents a binding site for RARA (Bowles et al., 2014; Marino et al., 2011), FOX (Niimi et al., 2017), SNAI2 (Hultgren et al., 2020) and JUN, which is part of the MAPK activation cascade downstream of VEGF signalling (reviewed in Guo et al., 2020) **(Fig. 2E, S3C, Table S4)**. -*14prox1a* is situated 14kbp upstream of *prox1a* and this enhancer is restricted to the facial lymphatics **(Fig. 2D)**. It also contains predicted binding sites for vascular relevant factors, such as ETS factors (Pham et al., 2007; De Val et al., 2008; Yoshimatsu et al., 2011), FOX TF (Niimi et al., 2017), TCFs (Cha et al., 2016; Nicenboim et al., 2015) and RARG (Bowles et al., 2014; Marino et al., 2011) (**Fig. 2E, S3D, Table S4-S5**).

In conclusion, we identified three additional lymphatic *prox1a* enhancers by means of chromatin state. Given that these elements show reduced or absent sequence conservation outside of Actinopterygii, this suggests evolutionary divergence in gene regulation – either through lineage-specific modification in *prox1a* regulation within ray-finned fish or a change at the level of DNA sequence while retaining function.

### -2.1prox1a_1 drives expression in the lymphatic valve

The -2.*1prox1a* element partially overlaps with that of a previously described -*11Prox1* murine lymphatic valve enhancer (Kazenwadel et al., 2023), specifically in its 5’ portion **(Fig. 3B)**. We hypothesised that this region might be sufficient to induce the valve-specific expression. The 200 bp fragment, called *-2.1prox1a_1*, was cloned in the ZED vector and tested *in vivo*. Expression driven by *-2.1prox1a_1* appeared at 3 dpf at the future valve position **(Fig. 3A, Fig. S4B-C)**, which mirrors the observations for the full enhancer. The enhancer driven valve expression was maintained at 5 dpf and 7 dpf **(Fig. 3A)**, aligning with the endogenous Prox1 protein distribution at 5 dpf, especially concentrated in the leaflets of the developing valves **(Fig. 3D)**. Similar to -2.*1prox1a*, -2.*1prox1a*_1 remained inactive in the trunk lymphatics **(Fig. 3A)** and all venous tissues **(Fig. 3A, Fig. S4F).** Furthermore, we explored the potential role for the 3’ portion of -2.*1prox1a*, here called *-2.1prox1a_2*, as a complementary contributor to -2.*1prox1a_1* activity. Upon *in vivo* testing, this element showed subtle GFP expression in the facial, but not in the trunk, lymphatics **(Fig. 3C, Fig. S4D)**. This suggests that this segment of

**Figure 3:**
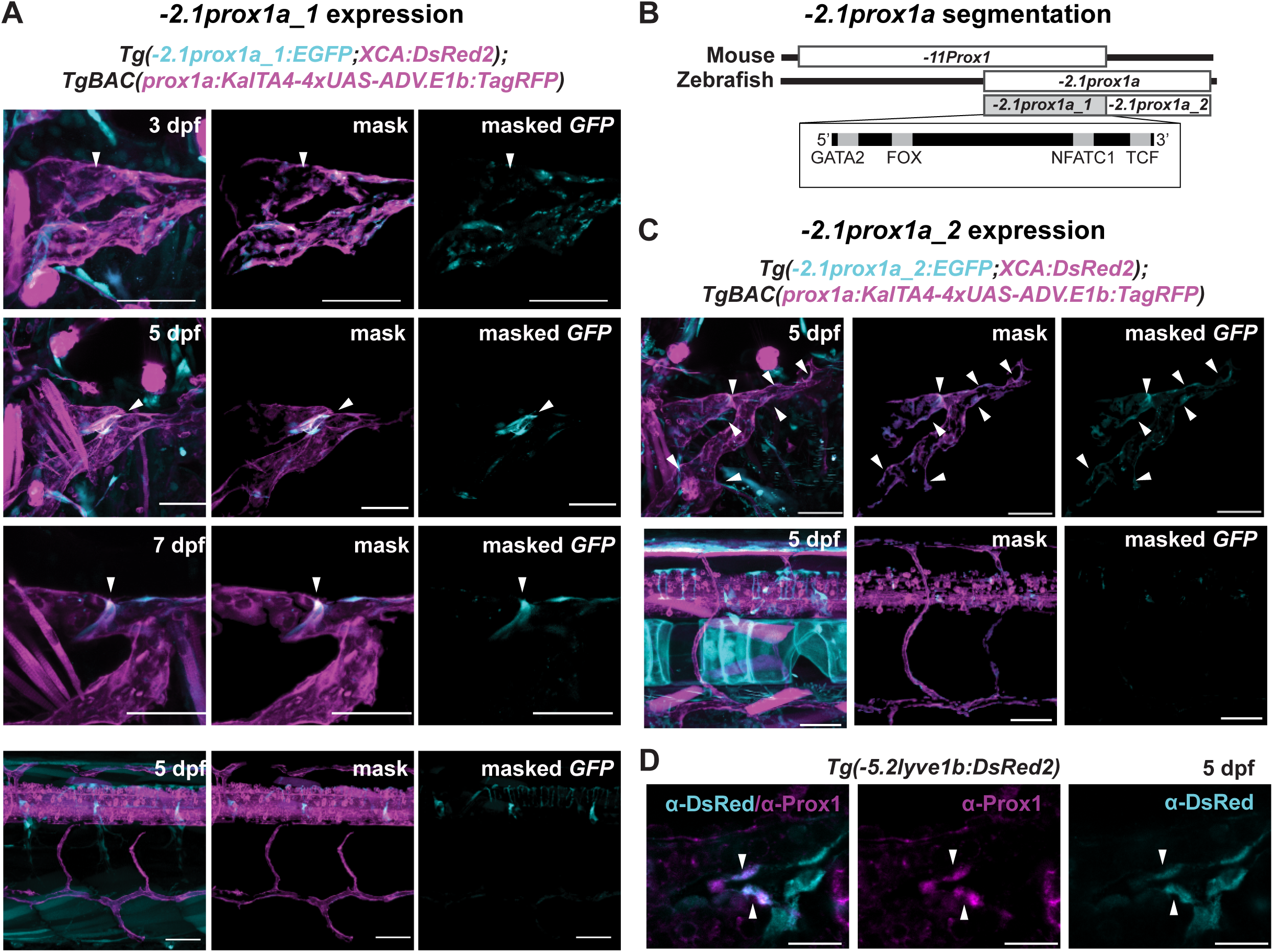
*-2.1prox1a_1* is the core element driving valve expression. (A) Confocal projections of the facial and trunk lymphatics labelled with *Tg(-2.1prox1a_1:EGFP; XCA:DsRed2)^uu5kk^* (cyan) and *Tg(prox1a:RFP)^nim5^* (magenta) at 3, 5 and 7 dpf. Arrows: Expression in the developing lymphatic valve. Scale bars: 50μm. (B) Schematic representation of the *-2.1prox1a* zebrafish and the *-11Prox1* mouse enhancer. The identified sequence overlap between the two enhancers is referred to as *-2.1prox1a_1* and the zebrafish unique enhancer part is referred to as *-2.1prox1a_2*. The TF binding sites location in *-2.1prox1a_1* is illustrated in the box. (C) Confocal projections of the facial and trunk lymphatics labelled with *Tg(-2.1prox1a_2:EGFP;XCA:DsRed2)^uu6kk^* (cyan) and *Tg(prox1a:RFP)^nim5^* (magenta) at 5 dpf. Arrowheads: Expression in the facial lymphatics. Scale bars: 50μm. (D) Confocal projections of the immunostaining against Prox1 (magenta) and *Tg(-5.2lyve1b:DsRed2)^nz101^* (cyan). Prox1 expression is detected in the valve leaflets at 5 dpf (arrows). Scale bar: 20μm.

### -2.1prox1a also plays a role in regulating prox1a expression

In summary, -2.*1prox1a*_1 serves as the core enhancer element of *-2.1prox1a* sufficient to drive expression in the developing valve from the early stages to later developmental phases.

### -2.1prox1a_1 is necessary for correct valve morphology and functionality

To test the necessity of these enhancers for the development of lymphatic vasculature, we concentrated on the core valve element *-2.1prox1a_1* and used CRISPR/Cas9 to generate the *en-2.1prox1a^uu12kk^* mutant line, hereafter referred to as Δ*-2.1prox1a,* carrying a 102 bp deletion covering the predicted NFATC1 binding site in the *-2.1prox1a_1* sequence. Homozygous mutants are viable, fertile and show normal body morphology (data not shown). We focused on the valve area to characterise the phenotype. Immunostainings revealed reduced Prox1a protein levels within the valve of mutant Δ*-2.1prox1a* compared to siblings (**Fig. 4A**), suggesting the deletion of the enhancer negatively impacts the regulation of *prox1a*. We then investigated gross vessel morphology at the valve position, which showed high resemblance between Δ*-2.1prox1a* homozygous and sibling embryos **(Fig. 4B-C-D-E)**. Similarly, the volume of the FCLV remained unaffected in Δ*-2.1prox1a* embryos **(Fig. S5A-B-C)**. Utilizing the average phenotype approach (Arnold et al., 2022) at 5 and 7 dpf, we further investigated the vessel morphology. No discernible differences were observed **(Fig. S5D)**, further confirming that the facial lymphatic vessels form accurately in the mutants. Conversely, an analysis of gross valve morphology at 7 and 14 dpf revealed an noticeable trend of leaflet alteration in the mutants compared to the sibling **(Fig. 4F).** Since 7 dpf is the stage in which the valve leaflets are completely formed (Shin et al., 2019), we further investigated the fine morphology of the valve using transmission electron microscopy (TEM). Imaging of three embryos per genotype revealed altered valve morphology in 7 dpf mutant embryos, with a reduction in length of both the cells composing the leaflet and the leaflets themselves **(Fig. 4G)**. Roundness of the valve cell nuclei was unaffected **(Fig. S5E)**.

**Figure 4:**
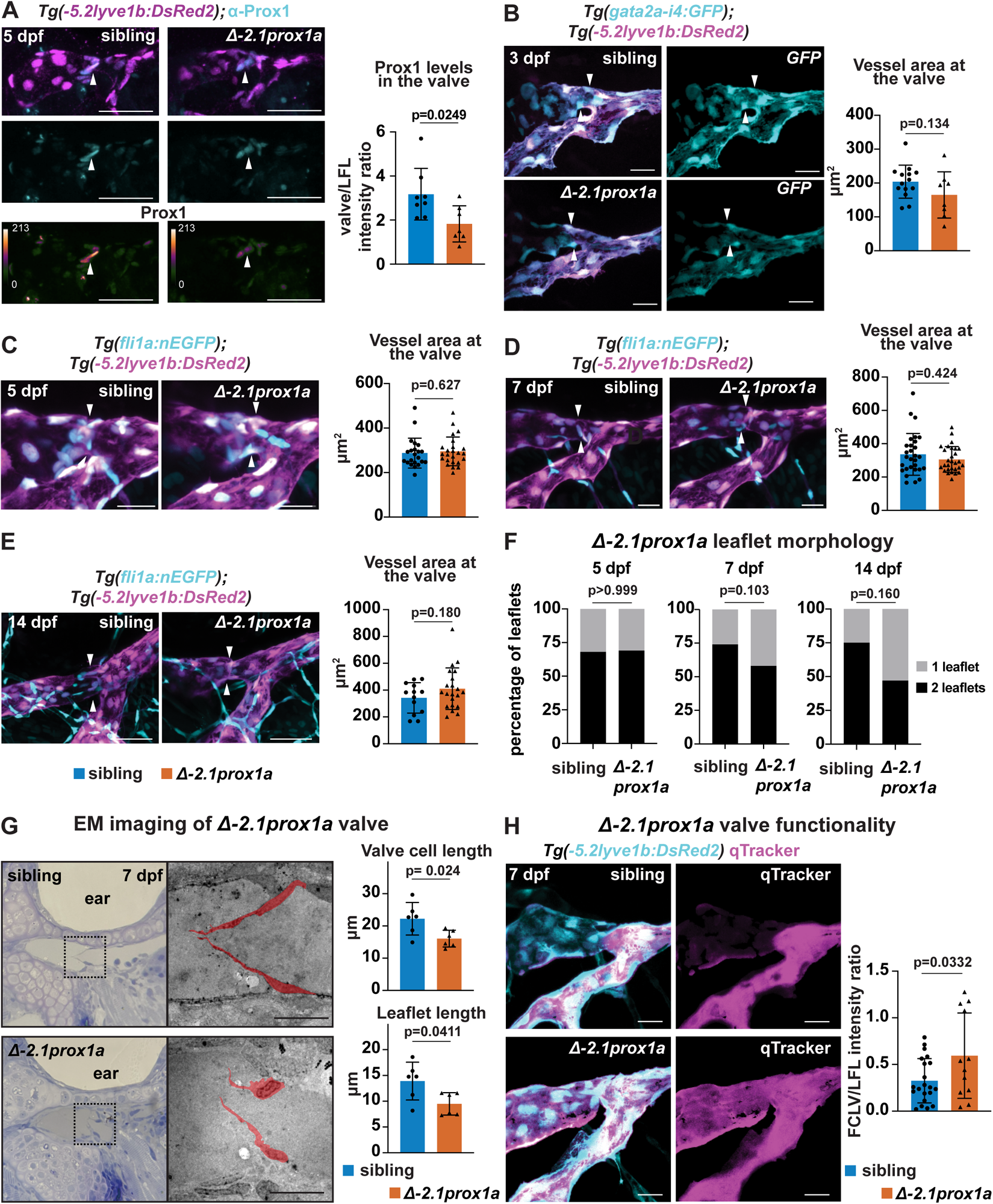
*-2.1prox1a* is necessary for correct valve development and function. (A) Top and middle: Confocal projections of immunostaining against Prox1 (cyan) and *Tg(-5.2lyve1b:DsRed2)^nz101^* (magenta) in sibling and 1′*-2.1prox1a* mutant embryos at 5 dpf. Bottom: Heatmap visualisation of Prox1 protein levels in sibling and 1′*-2.1prox1a* mutant embryos. Left: Quantification of Prox1 protein levels in the valves. Scale bar: 50μm. Sibling (n=8) vs mutants (n=7): Student’s t-test, p=0.0249. (B) Confocal projections of *Tg(gata-i4:GFP)^uu11kk^ (cyan)* and *Tg(-5.2lyve1b:DsRed2)^nz101^* (magenta) in sibling and 1′*-2.1prox1a* mutant embryos at 3 dpf. Quantification of vessel section at the valve (arrows) in 1′*-2.1prox1a* embryos at 3 dpf. Scale bar: 20μm. Sibling (n=14) vs mutants (n=8): Student’s t-test, ns (p=0.134). (C) Confocal projections of *Tg(fli1:nEGFP)^y7^ (cyan)* and *Tg(-5.2lyve1b:DsRed2)^nz101^* (magenta) in sibling and 1′*-2.1prox1a* mutant embryos at 5 dpf. Quantification of vessel section at the valve (arrows) in 1′*-2.1prox1a* embryos at 5 dpf. Scale bar: 20μm. Sibling (n=22) vs mutants (n=24), Mann-Whitney test, ns (p=0.627). (D) Confocal projections of *Tg(fli1:nEGFP)^y7^ (cyan)* and *Tg(-5.2lyve1b:DsRed2)^nz101^* (magenta) in sibling and 1′*-2.1prox1a* mutant embryos at 7 dpf. Quantification of vessel section at the valve (arrows) in 1′*-2.1prox1a* embryos at 7 dpf. Scale bar: 20μm. Sibling (n=29) vs mutants (n=29), Mann-Whitney test, ns (p=0.424). (E) Confocal projections of *Tg(fli1:nEGFP)^y7^ (cyan)* and *Tg(-5.2lyve1b:DsRed2)^nz101^* (magenta) in sibling and 1′*-2.1prox1a* mutant embryos at 14 dpf. Quantification of vessel section at the valve (arrows) in 1′*-2.1prox1a* embryos at 14 dpf. Scale bar: 50μm. Sibling (n=13) vs mutants (n=21): Student’s t-test, ns (p=0.180). (F) Quantifications of leaflet morphology in the valves of 5, 7 and 14 dpf sibling and 1′*-2.1prox1a* embryos. Fisher’s test of 5 dpf siblings (n=19) vs 1′*-2.1prox1a* (n=26), ns (p>0.999). Fisher’s test of 7 dpf siblings (n=27) vs 1′*-2.1prox1a* (n=29), ns (p=0.103). Fisher’s test of 14 dpf siblings (n=12) vs 1′*-2.1prox1a* (n=21), ns (p=0.160). (G) Brightfield and EM imaging of sibling (n=3) and 1′*-2.1prox1a* (n=3) valves at 7 dpf. The leaflets in the EM images are highlighted in red. Quantification of cell length: Student’s t-test, p= 0.024. Quantification of leaflet length: Student’s t-test, p= 0.0411. Scale bar: 10μm. (H) Confocal projections of *Tg(-5.2lyve1b:DsRed2)^nz101^ (cyan)* and qTracker (magenta) in sibling and 1′*-2.1prox1a* mutant embryos at 7 dpf, visualising flow through the vessel. Quantification of Qtracker leakage through the valve in sibling and 1′*-2.1prox1a* embryos at 7 dpf. Scale bar: 20μm. Siblings (n=21) vs mutants (n=12), Student’s t-test, p=0.0332.

To assess whether the altered morphology could affect valve function, we performed Qtracker injections in the LFL in anaesthetised 7 dpf larvae as previously described (Shin et al., 2019). Leakage through the valve was scored and quantified. Our findings demonstrated increased leakage in ι1*-2.1prox1a* mutants compared to siblings, confirming a functional impairment of the lymphatic valve in the absence of the enhancer **(Fig. 4H, S5F)**. Collectively, these data show the necessity for *-2.1prox1a_1* for correct valve formation and proper expression of *prox1a*, highlighting the functional importance of *prox1a* enhancers for precise lymphatic development.

## Discussion

PROX1 is a key factor for LEC development (Koltowska et al., 2015a; Wigle and Oliver, 1999). Despite this, only one *cis*-regulatory element of *Prox1* have been identified so far (Kazenwadel et al., 2023). Here, we used a mixed approach, taking advantage of both evolutionary conservation and tissue-specific chromatin accessibility, to identify *prox1a* enhancers active in the lymphatic endothelium of zebrafish. Unexpectedly, we could identify five separate enhancers driving reporter expression in the lymphatic vasculature. While two of the enhancers, *-2.1prox1a* and *+15.2prox1a,* show significantly enriched expression in anatomically and functionally distinct subsets of the lymphatics, such as the FCLV and the valve, the activity of the three remaining enhancers overlaps to a larger extent. Specifically, both *-87prox1a* and *-71prox1a* drive expression in trunk and facial lymphatics. Enhancers with high redundancy in their expression patterns are defined as shadow enhancers (Hobert, 2010; Hong et al., 2008). Shadow enhancers are a common feature among developmental genes (Cannavò et al., 2016; Kvon et al., 2021). They are thought to serve as a mechanism aimed at ensuring robustness during the developmental process (Antosova et al., 2016; Kvon et al., 2021; Osterwalder et al., 2018). Various types of interactions - additive, superadditive, subadditive, and repressive - among these type of enhancers have been shown to effectively fine-tune the regulation of target genes into specific patterns (Bothma et al., 2015; El-Sherif and Levine, 2016; Kvon et al., 2021; Lam et al., 2015). Moreover, studies indicate that shadow enhancers can form transcriptional hubs, where multiple enhancers concurrently interact with the gene promoter at the same time (Kvon et al., 2021). Given this intricate interplay, the presence of two enhancers, *-87prox1a* and *-71prox1a*, with extensively overlapping expression suggest potential role for shadow enhancers in *prox1a* regulation. The complex regulatory logic at *prox1a* locus highlights the necessity for tight regulation of *prox1a* activity to ensure a correct lymphatic development.

Our conservation analysis revealed high density of conserved non-coding elements (CNEs) surrounding the *prox1a* locus. As CNEs can be considered putative enhancers, this suggests *cis*-regulation plays an important role in controlling *prox1a* expression in more than just LECs. In fact, among 10 tested CNEs, only two functional lymphatic enhancers were identified: *- 2.1prox1a* and *+15.2prox1a*. In contrast, three sequences identified by chromatin state, *- 87prox1a*, *-71prox1a* and *-14prox1a,* presented varying conservation levels within Actinopterygii. Although distal enhancers have a tendency to be less conserved, the presence of other vertebrate-conserved CNEs spanning the 150 kilo base pairs (kbp) upstream region of *prox1a* suggests no major rearrangement have contributed to the loss of these enhancers in tetrapods. In addition, we observed conservation of microsynteny surrounding *prox1a* in tetrapods and actinopterygians, with the exception of an Otocephala specific divergence. Such conservation in loci disposition also speaks against major genomic rearrangements in the region. Interestingly, the most conserved enhancers at the sequence level also have the most spatially restricted expression, respectively the lymphatic valve and the FCLV, while the less conserved enhancer sequences with tetrapods display broader and overlapping expression patterns. The majority of the binding sites for well described lymphatic regulators in mammals, such as GATA2, PROX1 and NFATC1, are found in the sequence conserved enhancer. This suggests potential divergence in the molecular code upstream of *-87prox1a*, *-71prox1a* and *- 14prox1a* between Actinopterygii and Tetrapoda, potentially involving different transcription factors. However, it is important to note that evolutionary conservation of functional enhancers does not necessarily require high levels of conservation at a sequence level, as long as key transcription factor motifs endure (Wong et al., 2020). As the nature of the enhancers can be complex, adopting complementary methods for their identification is crucial for a comprehensive understanding of a gene regulatory landscape.

Sequence conservation can serve as an indicator of enhancer presence; however, it does not guarantee complete functional conservation. In the case of *-11Prox1/-2.1prox1a*, the element drives expression in the lymphatic valve in both mouse and zebrafish. Nevertheless, profound differences in the mutant phenotypes are reported. *-11Prox1* mutants have more severe lymphatic defects and die perinatally (Kazenwadel et al., 2023). Conversely, the *-2.1prox1a* mutants only show developmental defects in the lymphatic valve, and are viable and fertile. The deletion of *-11Prox1* is fully phenocopied by the deletion of the GATA2 binding site. This GATA2 site is responsible for the transition of LECs to HECs in *-11Prox1* mutant mice (Kazenwadel et al., 2023). However, the putative GATA2 binding site is not conserved in zebrafish *-2.1 prox1a* and therefore the hematopoietic phenotypes were not further explored in this study. The observed differences between *-11Prox1* and *-2.1prox1a* suggest functional divergence despite their common evolutionary origin. This highlights that that functional conservation cannot be solely deduced from enhancer sequence data alone.

In conclusion, this study has demonstrated that *prox1a* in the lymphatic vasculature is regulated by a diverse group of distinct enhancers. We uncovered that the distal enhancers drive the expression in large regions of the lymphatic endothelium, whereas the proximal and sequence conserved enhancers channel the expression to the functionally distinct sub-compartments of the developing lymphatic vascular network. This work represents a first step towards a full characterisation of the cis-regulatory landscape of *prox1a* and the understanding of the complex mechanisms regulating its lymphatic expression.

## Acknowledgements

This work was supported by Wallenberg Academy Fellowship (2017.0144), Ragnar Söderbergs Fellowship (M13/17), Vetenskapsådet (VR-MH-2016-01437) and Beijer Foundation. The Genome Engineering Zebrafish at SciLifeLab provided the *en-2.1 prox1a_1* mutant line. Electron microscopy was performed at BioVis at Uppsala University.

## Material and Methods

### Zebrafish

Zebrafish work was carried out with ethical approval from the Swedish Board of Agriculture (5.8.18-10590/2018) and in compliance with the animal ethics committees at the Peter MacCallum Cancer Centre, The University of Melbourne. The fish were maintained at the Genome Engineering Zebrafish National Facility (SciLifeLab, Uppsala Sweden) and the Danio Rerio University of Melbourne facility (DrUM, Melbourne, Australia). Adults and embryos were housed according to standard procedures. The previously published lines used in this study were *Tg(-5.2lyve1b:DsRed2)^nz101^*(Okuda et al., 2012)*, TgBAC(prox1a:KalTA4-4xUAS-ADV.E1b:TagRFP)^nim5^,* refer to as *Tg(prox1a:RFP)^nim5^* in this study, (Dunworth et al., 2014; van Impel et al., 2014), *Tg(kdr-l:ras-cherry)^s916^* (Hogan et al., 2009), *Tg(fli1a:nEGFP)^y7^*(Lawson and Weinstein, 2002) and *Tg(-2.1prox1a:EGFP;XCA:DsRed2)^uu3kk^*(Kazenwadel et al., 2023).

The *Tg(-2.1prox1a_1:EGFP;XCA:DsRed2)^uu5kk^, Tg(-2.1prox1a_2:EGFP; XCA:DsRed2)^uu6kk^*, *Tg(+15.2prox1a:EGFP;XCA:DsRed2)^uu7kk^, Tg(-2.1prox1a_3:EGFP; XCA:DsRed2)^uom119^, Tg(-71prox1a:EGFP; XCA:DsRed2)^uom121^, Tg(-87prox1a:EGFP; XCA:DsRed2)^uom122^, Tg(-14prox1a:EGFP; XCA:DsRed2)^uom120^ Tg(-2.1prox1a:basEGFP;ACry:GFP)^uu10kk^*, *en.-2.1prox1a^uu12kk^* and *Tg(gata-i4:GFP)^uu11kk^* lines were generated for this study.

### In-silico predictions

To identify of sequence-conserved enhancers, the DNA regions between the two loci neighbouring *Prox1*, *prox1a,* or *prox3* were downloaded from ENSEMBL, together with annotations. For *prox3*, homologs were identified by BLAST of zebrafish *prox3* and confirmed by local synteny. The sequences were oriented in the direction of the transcription. The species, assemblies and regions used are listed in **Table S1**. Conserved non-coding elements were identified using mVista non-coding DNA conservation analysis (Dubchak et al., 2000; Frazer et al., 2004; Mayor et al., 2000). The alignment was performed using the LAGAN program (Brudno et al., 2003).

To determine the zebrafish-specific binding sites in the identified *prox1a* enhancers, we conducted a motif discovery analysis of zebrafish enhancer sequences using FIMO in the MEME Suite (Grant et al., 2011). Our motif prediction was constrained to a search for motifs of size 7, with a p-value cut-off of 1e-04. A comprehensive list of predicted motifs is shown in **Table S6**.

To identify evolutionarily conserved motifs and binding sites within the specified enhancers, we retrieved the identified conservation peaks and subjected them to analysis using the MEME Suite (Bailey and Elkan, 1994) and TOMTOM (Gupta et al., 2007). To validate the predicted conserved binding sites, we compared the results of this analysis with the outcomes of the FIMO motif discovery analysis, utilising a p-value cut-off of 1e-02. The decision to use a higher p-value stems from the conservation of these sequences and aims to capture an informative overview of the potential binding sites. Binding sites for known endothelial factors that appear in both analyses have been reported.

### snATAC-seq data processing and analysis

For snATAC-seq data analysis, we included only the 4 dpf wild type cells from the publicly available data set (Grimm et al., 2023). The analysis was performed and data were processed as previously described (Grimm et al., 2023).

All gene ontology analyses were performed using Panther.db (Thomas et al., 2006) (Biological Process Complete). The complete list of the predicted GO terms is shown in **Table S7**.

### Cloning and transgenesis

The conserved elements of interest (**Table S2 and S3)** were cloned into the ZED vector as previously described (Bessa et al., 2009). The *Tg(-2.1prox1a:basEGFP;ACry:GFP)^uu10kk^*line was generated by cloning the *-2.1prox1a* sequence in a p5E-MCS vector (Quillien et al., 2017) (Addgene #26029) using In-Fusion cloning (Takara Bio, primers are listed in **Table S3**) and then inserted into a pDestTol2ACryGFP backbone (Berger and Currie, 2013) (Addgene #64022), with a pENTRbasEGFP (Villefranc et al., 2007) (Addgene #22453) and a p3E-polyA (Tol2kit v1.2 #302) vector using the Gateway cloning method (Invitrogen). ATAC-identified enhancers were inserted into the ZED vector by In-Fusion cloning (Cat. #638910, In-Fusion HD Cloning Plus Kits, Takara Bio) using BspEI and BmgBI cutting sites for linearisation. Primers used are listed in **Table S3**. To generate transgenic lines, 1ul of construct at 20 ng/μl and tol2 transposase mRNA at 100 ng/μl were injected into the one-cell stage wild type zebrafish embryos. F0 embryos were screened for reporter expression and F1 embryos were screened using confocal microscopy for GFP expression in the lymphatic structures. The numbers of F0 founders screened for each tested CRE are listed in **Table S2**. Bleed-through in the GFP channel was excluded by imaging the embryos that were negative for *TgBAC(prox1a:KalTA4-4xUAS-ADV.E1b:TagRFP)^nim5^* **(Fig. S4E)**. F1 fish with lymphatic GFP expression were used to establish the stable lines. The *Tg(gata2a-i4-1.1 kb:GFP)^uu11kk^* was created by injecting the construct (74) as previously described.

### Imaging and image processing

For the conserved enhancer reporter and mutant lines, transgenic embryos were anesthetised with tricaine, mounted in 1% low-melting agarose, and face or trunk was imaged using a Leica TCS SP8 DLS microscope with a Fluotar VISR 25X water objective (objective number:11506375). For enhancers identified by snATAC-seq, transgenic embryos were anesthetised with tricaine, mounted in 0.5% low-melting agarose and imaging was conducted at the Centre for Advanced Histology and Microscopy (Peter MacCallum Cancer Centre). Live samples were imaged using a Zeiss LSM 780 FCS confocal microscope. Images were processed using ImageJ software version 2.9.0. Immunostained embryos were mounted in clearing solution Omnipaque (350 mg litre^−1^ concentration per 1 ml iohexol, GE Healthcare) and imaged using Leica TCS SP8 DLS microscope as described above. Masking was performed in Imaris v9.3.0 with a surface detail of 1μm. All representative images are Maximum Intensity Projections of the z-stack generated using ImageJ 2.9.0. The skin signals in the GFP channel were manually removed.

### Mutant line generation

Mutants were generated using CRISPR/Cas9 as described previously (Carrington et al., 2015; Varshney et al., 2016). The guides were designed to flank the target enhancer sequences. Zebrafish embryos were injected at one-cell stage with the 70–140 ng/μL of each gRNA and 200 ng/μL Cas9 mRNA. These guides are listed in **Table S8.** Fish were genotyped by PCR, followed by gel electrophoresis, and the deletion was confirmed by Sanger sequencing (Eurofins). F1 embryos with identical mutations were used to establish the stable mutant line.

### Genotyping

Δ*-2.1prox1a* embryos were genotyped by PCR, followed by gel electrophoresis. PCR was performed as previously described for FLA genotyping (Carrington et al., 2015) using the primers listed in **Table S3**. Fluorescent primers were omitted from the reaction.

### Immunostaining

Immunostaining was performed as previously described (Le Guen et al., 2014; Shin et al., 2016), with the addition of a 45 min RT digestion step in PK as described previously (Koltowska et al., 2015b). The primary antibodies used were α-Prox1 rabbit (AngioBio #11-002P) and α-mCherry chicken (AvesLabs #MCHERRY-0020). The secondary antibodies used were α-rabbit IgG HRP (Cell Signalling Technology #7076) and α-Chicken 488 (Jackson Immuno Leb #703-545-155). Signal amplification was performed using the TSA^TM^ Plus Cyanine 3 System (Perkin Elmer #NEL744001KT), with a development time of 3 h. Imaging was performed as described above.

### Electron Microscopy imaging

For transmission electron microscopy (TEM), Δ*-2.1prox1a* embryos were fixed in 2.5% Glutaraldehyde (Ted Pella) + 1% Paraformaldehyde (Merck) in 0.1 M Phosphate buffer (PB) pH 7.4, then embedded in 8% agar and a 300μm sagittal section cut on a Microm M 650V vibratome (ThermoScientific) and stored at 4°C until further processed. The tails were then used for genotyping. Samples were rinsed with 0.1 M PB for 10 min prior to 1 h incubation in 1% osmium tetroxide (TAAB) in 0.1 M PB. After rinsing in 0.1 M PB, samples were dehydrated using increasing concentrations of ethanol (50%, 70%, 95% and 99.9%) for 10 minutes each step, followed by 5 min incubation in propylene oxide (TAAB). The samples were then placed in a mixture of Epon Resin (Ted Pella) and propylene oxide (1:1) for 1 h, followed by 100% resin and left o/n. Subsequently, samples were embedded in capsules in newly prepared Epon resin and left for 1-2 h and then polymerized at 60°C for 48 h.

Semi-thin sections were cut, stained with Toluidine blue, and examined using light-microscopy to identify the area of interest. Ultrathin sections (60-70 nm) were cut in an EM UC7 Ultramicrotome (Leica) and placed on a grid. The sections were subsequently contrasted with 5% uranyl acetate and Reynold’s lead citrate and visualised with Tecnai™ G2 Spirit BioTwin transmission electron microscope (Thermo Fisher/FEI) at 80 kV with an ORIUS SC200 CCD camera and Gatan Digital Micrograph software (both from Gatan Inc.).

### Qtracker injections

Microangiography was performed following a previously published protocol (Arnold et al., 2022; Shin et al., 2019). Embryos were anaesthetised with Tricane and injected with 1nL of Qtracker™ 655 Vascular labels (Thermo Fisher) at 7 dpf in the LFL for valve leakage experiments and imaged on a Leica TCS SP8 DLS microscope immediately post-injection, approximately 5 min post-injection.

### Image quantification

The relative levels of Prox1 protein in the valve were calculated as the ratio between the representative valve and LFL nucleus. Briefly, the chosen nuclei were masked in Imaris v9.3.0 as described above. The average intensity was calculated and used to compile the ratio. Nuclei with a maximum intensity of 255 were excluded from the calculations.

Vessel sections were calculated in ImageJ 2.9.0 using a custom-made script. Briefly, the z-stack was rotated along its main axis to ensure that the vessel section was perpendicular to the cutting planes. The stack was then re-sliced along the yz-plane, and the valve position was selected. Thresholding was used to mask of the vessel area which was subsequently measured.

The FCLV volume was calculated in Imaris v9.3.0, by manually creating a surface spanning the 400 pixels of FCLV past the valve position and calculating its volume.

To generate average phenotypes, images were acquired and processed as previously described (Arnold et al., 2022).

Valve leaflet morphology was scored in Imaris v9.3.0, looking at the projections of the stack in x, y, and z, and scoring whether one or two leaflets were visible in any of them.

To calculate the valve cell length in the EM images, the cells composing the leaflets were traced manually and used to create a mask in ImageJ. The mask was then skeletonized and secondary branches were removed to obtain the length of the cell. Nuclei roundness was calculated in ImageJ by the “Round” image descriptor. Leaflet length was calculated by tracing the internal edge of the leaflet in ImageJ and measured.

Valve leakage in the injected embryos was quantified as the Qtracker signal ratio between the FCLV and LFL in a single slice.

### Statistical analysis

The normality of all numerical datasets was tested with a Shapiro-Wilk test. For pair-wise comparison, an unpaired two-tailed Student’s t-test was run on normally distributed data, while a Mann-Whitney test was run if normality was not confirmed. For frequency data, Fisher’s exact test was used.

**Figure S1:**

(A) Microsynteny surrounding the vertebrate *Prox1/prox1a* locus. Downstream of *Prox1/prox1a*, *Smyd2* is found in all analysed sequences. Upstream of *Prox1/prox1a, Rps6ck1* is found in all species, except zebrafish. The annotations used are listed in Table S1.

(B) Conservation analysis of the 54 kbp region surrounding the *prox3* locus comparted to nine vertebrate species. Results are shown compared to zebrafish. Blue peaks: Exons; red peaks: Region of conserved non-coding DNA; black arrowhead: Conserved peaks.

(C) H3K4me1 and H3K27ac at 48 hpf in whole zebrafish embryos (published dataset from Bogdanovic et al, 2012). Histone marks enrichment for the -2.*1prox1a* and +15.*2prox1a* sequences.

(D) Confocal projections of trunk lymphatics labelled with *Tg(+15.2prox1a:EGFP; XCA:DsRed2)^uu7kk^* (cyan) and *Tg(prox1a:RFP)^nim5^* (magenta) at 5 dpf. Arrows: Lymphatic vessels. Asterisks: Posterior cardinal vein. Scale bars: 50μm.

(E) Confocal projections of facial (top) and trunk (bottom) lymphatics at 5 dpf labelled with the empty ZED vector as a control. Arrows: Neuronal structures. Scale bars: 50μm.

(F) Confocal projections of facial and trunk lymphatics at 5 dpf labelled with *Tg(prox1a:RFP)^nim5^* (magenta) and injected with *empty:basEGFP;ACry:GFP* plasmid to create a transient control. Scale bars: 50μm.

(G) Conserved motif structure of the identified lymphatic *+15.2prox1a* enhancer. The motif key indicates the motive numbers corresponding to the colours and the motif sequences are listed in Table S4.

(H) Conserved motif structure of the identified lymphatic -2.*1prox1a* enhancer. The motif sequences are listed in Table S4.

(I) Predicted conserved binding site for *+15.2prox1a* in vertebrates. Motif locations are listed in Table S4.

(J) Predicted conserved binding site for -*2*.*1prox1a* in vertebrates. Motif locations are listed in Table S4.

**Figure S2:**

(A) Clusters of endothelial cells (ECs) and endocardium at 4 dpf with umaps showing the chromatin state at known lymphatics and venous loci. High accessibility at *prox1a* and *cdh6* marks the lymphatic endothelial cell (LECs) cluster. High *kdrl* accessibility marks the venous endothelial cells (VECs) and arterial endothelial cell (AECs) cluster, and high *lyve1b* accessibility marks LECs and VECs. muLECs: Mural lymphatic endothelial cells.

(B) GO enrichment of developmentally relevant terms in differentially accessible peaks in LECs.

(C) Schematic representation of the *-2.1prox1a* enhancer indicating the overlap between the *-2.1prox1a_3* sequence from the identified peak using the ATAC-sequencing, *-2.1prox1a* identified using the conservation method (Figure 1) and the segmentation into *-2.1prox1a*_1 and *-2.1prox1a*_2 (Figure 3). Predicted binding for lymphatic-associated transcription factors is indicated below the enhancer fragments. Blue: Binding sites identified in zebrafish (p-val < 1e-04). Red: Conserved binding sites within vertebrates. Yellow: Binding sites conserved in vertebrates but absent in zebrafish.

(D) Confocal projections of facial lymphatics labelled with *Tg(-2.1prox1a_3:EGFP; XCA:DsRed2)^uom119^* (cyan) and *Tg(prox1a:RFP)^nim5^* (magenta) at 5 dpf. Arrows: Expression in the developing lymphatic valve. Asterisks: expression in the facial lymphatic endothelium. Scale bars: 50μm.

**Figure S3:**

(A) The microsynteny surrounding the Actinopterygii *prox1a* locus. Downstream of *prox1a*, *Smyd2a* is found in all analysed sequences. Upstream of *Prox1/prox1a, Rps6ck1* is found in *Lepisosteus oculatus* and Euteleostei, but not in Otocephala. The annotations used are listed in Table 1.

(B) Conserved motif structure and predicted conserved binding sites of the identified lymphatic *-87prox1a* enhancer in Actinopterygii. Motif locations are listed in Table S4 and S5.

(C) Conserved motif structure and predicted conserved binding sites of the identified lymphatic *-71prox1a* enhancer in Actinopterygii. Motif locations are listed in Table S4 and S5.

(D) Conserved motif structure and predicted conserved binding sites of the identified lymphatic *-14prox1a* enhancer in Actinopterygii. Motif locations are listed in Table S4 and S5.

**Figure S4:**

(A) Confocal projections of VA-L labelled with *Tg(+15.2prox1a:EGFP;XCA:DsRed2)^uu7kk^* (cyan) and *Tg(prox1a:RFP)^nim5^*(magenta) at 54 hpf, showing no lymphatic expression.

(B) Confocal projections of facial lymphatics labelled with *Tg(-2.1prox1a_1:EGFP; XCA:DsRed2)^uu5kk^* (cyan) and *Tg(prox1a:RFP)^nim5^* (magenta) at 54 hpf, showing no lymphatic expression.

(C) Confocal projections of VA-L labelled with *Tg(-2.1prox1a_1:EGFP;XCA:DsRed2)^uu5kk^* (cyan) and *Tg(kdr-l:ras-cherry)^s916^*(magenta) at 54 hpf, showing no lymphatic expression.

(D) Confocal projections of facial lymphatics labelled with *Tg(-2.1prox1a_2:EGFP;XCA:DsRed2)^uu6kk^* (cyan) and *Tg(kdr-l:ras-cherry)^s916^* (magenta) at 54 hpf, showing -2.*1prox1a_2* driven expression in the VA-L (arrow).

(E) Enhancer-driven expression in *Tg(-2.1prox1a_1:EGFP;XCA:DsRed2)^uu5kk^*, *Tg(-2.1prox1a_2:EGFP;XCA:DsRed2)^uu6kk^* and *Tg(+15.2prox1a:EGFP;XCA:DsRed2)^uu7k^*in the facial lymphatics (arrows) at 5 dpf, excluding bleed-through as a source of the signal.

(F) Confocal projections of the primary head sinus labelled with *Tg(-2.1prox1a_1:EGFP;XCA:DsRed2)^uu5kk^*, *Tg(-2.1prox1a_2:EGFP;XCA:DsRed2)^uu6kk^*and *Tg(+15.2prox1a:EGFP;XCA:DsRed2)^uu7k^* at 5 dpf, showing +*15.2prox1a* driven venous expression (arrow).

**Figure S5:**

(A-C): Quantification of FCLV volume in 1′*-2.1prox1a* embryos at 5 (A), 7 (B) and 14 (C) dpf.

(A) Student’s t-test of 5 dpf siblings (n=16) vs 1′*-2.1prox1a* (n=22), ns (p=0.064). (B) Student’s t-test of 7 dpf siblings (n=21) vs 1′*-2.1prox1a* (n=26), ns (p=0.236). (C) Student’s t-test of 14dpf siblings (n=8) vs 1′*-2.1prox1a* (n=17), ns (p=0.187).

(D) Average valve phenotype in sibling (cyan) vs 1′*-2.1prox1a* (magenta) embryos at 5 dpf and 7 dpf, showing similar gross vessel morphology. Scale bar: 20μm.

(E) Quantification of sibling (n=6) vs 1′*-2.1prox1a* mutant (n=6) valve cell nuclei roundness in TEM-imaged embryos at 7 dpf, representative images are present in Figure 4G. Student’s t-test, ns (p=0.286).

(F) Scoring of valve leakage in Qtracker injected 7 dpf siblings (n=21) vs 1′*-2.1prox1a* (n=12) embryos, representative image are present in Figure 4H. Fisher’s test, p=0.026.

## Notes

### Competing Interest Statement

The authors have declared no competing interest.

## Bibliography

Aday, A.W., Zhu, L.J., Lakshmanan, A., Wang, J., and Lawson, N.D. (2011). Identification of cis regulatory features in the embryonic zebrafish genome through large-scale profiling of H3K4me1 and H3K4me3 binding sites. Dev. Biol. 357, 450–462.

Antosova, B., Smolikova, J., Klimova, L., Lachova, J., Bendova, M., Kozmikova, I., Machon, O., and Kozmik, Z. (2016). The Gene Regulatory Network of Lens Induction Is Wired through Meis-Dependent Shadow Enhancers of Pax6. PLOS Genet. 12, e1006441.

Arnold, H., Panara, V., Hußmann, M., Filipek-Gorniok, B., Skoczylas, R., Ranefall, P., Gloger, M., Allalou, A., Hogan, B.M., Schulte-Merker, S., et al. (2022). mafba and mafbb differentially regulate lymphatic endothelial cell migration in topographically distinct manners. Cell Rep. 39, 110982.

Bailey, T.L., and Elkan, C. (1994). Fitting a Mixture Model by Expectation Maximization to Discover Motifs in Bipolymers. Proc. Second Int. Conf. Intell. Syst. Mol. Biol. 28–36.

Berger, J., and Currie, P.D. (2013). 503unc, a small and muscle-specific zebrafish promoter. Genesis 51, 443–447.

Bessa, J., Tena, J.J., de la Calle-Mustienes, E., Fernández-Miñán, A., Naranjo, S., Fernández, A., Montoliu, L., Akalin, A., Lenhard, B., Casares, F., et al. (2009). Zebrafish Enhancer Detection (ZED) vector: A new tool to facilitate transgenesis and the functional analysis of cis-regulatory regions in zebrafish. Dev. Dyn. 238, 2409–2417.

Bogdanovic, O., Fernandez-Minan, A., Tena, J.J., de la Calle-Mustienes, E., Hidalgo, C., van Kruysbergen, I., van Heeringen, S.J., Veenstra, G.J.C., and Gomez-Skarmeta, J.L. (2012). Dynamics of enhancer chromatin signatures mark the transition from pluripotency to cell specification during embryogenesis. Genome Res. 22, 2043–2053.

Bonn, S., Zinzen, R.P., Girardot, C., Gustafson, E.H., Perez-Gonzalez, A., Delhomme, N., Ghavi-Helm, Y., Wilczyński, B., Riddell, A., and Furlong, E.E.M. (2012). Tissue-specific analysis of chromatin state identifies temporal signatures of enhancer activity during embryonic development. Nat. Genet. 44, 148–156.

Bothma, J.P., Garcia, H.G., Ng, S., Perry, M.W., Gregor, T., and Levine, M. (2015). Enhancer additivity and non-additivity are determined by enhancer strength in the Drosophila embryo. Elife 4.

Bowles, J., Secker, G., Nguyen, C., Kazenwadel, J., Truong, V., Frampton, E., Curtis, C., Skoczylas, R., Davidson, T.-L., Miura, N., et al. (2014). Control of retinoid levels by CYP26B1 is important for lymphatic vascular development in the mouse embryo. Dev. Biol. 386, 25–33.

Brudno, M., Do, C.B., Cooper, G.M., Kim, M.F., Davydov, E., Green, E.D., Sidow, A., and Batzoglou, S. (2003). LAGAN and Multi-LAGAN: Efficient tools for large-scale multiple alignment of genomic DNA. Genome Res.

Bussmann, J., Bos, F.L., Urasaki, A., Kawakami, K., Duckers, H.J., and Schulte-Merker, S. (2010). Arteries provide essential guidance cues for lymphatic endothelial cells in the zebrafish trunk. Development 137, 2653–2657.

Cannavò, E., Khoueiry, P., Garfield, D.A., Geeleher, P., Zichner, T., Gustafson, E.H., Ciglar, L., Korbel, J.O., and Furlong, E.E.M. (2016). Shadow Enhancers Are Pervasive Features of Developmental Regulatory Networks. Curr. Biol. 26, 38–51.

Carrington, B., Varshney, G.K., Burgess, S.M., and Sood, R. (2015). CRISPR-STAT: An easy and reliable PCR-based method to evaluate target-specific sgRNA activity. Nucleic Acids Res.

Cha, B., Geng, X., Mahamud, M.R., Fu, J., Mukherjee, A., Kim, Y., Jho, E., Kim, T.H., Kahn, M.L., Xia, L., et al. (2016). Mechanotransduction activates canonical Wnt/β-catenin signaling to promote lymphatic vascular patterning and the development of lymphatic and lymphovenous valves. Genes Dev. 30, 1454–1469.

Chiang, I.K.-N., Fritzsche, M., Pichol-Thievend, C., Neal, A., Holmes, K., Lagendijk, A., Overman, J., D’Angelo, D., Omini, A., Hermkens, D., et al. (2017). SoxF factors induce Notch1 expression via direct transcriptional regulation during early arterial development. Development 144, 2629–2639.

Creyghton, M.P., Cheng, A.W., Welstead, G.G., Kooistra, T., Carey, B.W., Steine, E.J., Hanna, J., Lodato, M.A., Frampton, G.M., Sharp, P.A., et al. (2010). Histone H3K27ac separates active from poised enhancers and predicts developmental state. Proc. Natl. Acad. Sci. 107, 21931–21936.

Dieterich, L.C., Klein, S., Mathelier, A., Sliwa-Primorac, A., Ma, Q., Hong, Y.-K., Shin, J.W., Hamada, M., Lizio, M., Itoh, M., et al. (2015). DeepCAGE Transcriptomics Reveal an Important Role of the Transcription Factor MAFB in the Lymphatic Endothelium. Cell Rep. 13, 1493–1504.

Dubchak, I., Brudno, M., Loots, G.G., Pachter, L., Mayor, C., Rubin, E.M., and Frazer, K.A. (2000). Active conservation of noncoding sequences revealed by three-way species comparisons. Genome Res.

Dunworth, W.P., Cardona-Costa, J., Bozkulak, E.C., Kim, J.-D., Meadows, S., Fischer, J.C., Wang, Y., Cleaver, O., Qyang, Y., Ober, E.A., et al. (2014). Bone Morphogenetic Protein 2 Signaling Negatively Modulates Lymphatic Development in Vertebrate Embryos. Circ. Res. 114, 56–66.

El-Sherif, E., and Levine, M. (2016). Shadow Enhancers Mediate Dynamic Shifts of Gap Gene Expression in the Drosophila Embryo. Curr. Biol. 26, 1164–1169.

Eng, T.C.Y., Chen, W., Okuda, K.S., Misa, J.P., Padberg, Y., Crosier, K.E., Crosier, P.S., Hall, C.J., Schulte-Merker, S., Hogan, B.M., et al. (2019). Zebrafish facial lymphatics develop through sequential addition of venous and non-venous progenitors. EMBO Rep. 20, 1–17.

François, M., Caprini, A., Hosking, B., Orsenigo, F., Wilhelm, D., Browne, C., Paavonen, K., Karnezis, T., Shayan, R., Downes, M., et al. (2008). Sox18 induces development of the lymphatic vasculature in mice. Nature 456, 643–647.

Frazer, K.A., Pachter, L., Poliakov, A., Rubin, E.M., and Dubchak, I. (2004). VISTA: Computational tools for comparative genomics. Nucleic Acids Res. 32.

Frye, M., Taddei, A., Dierkes, C., Martinez-Corral, I., Fielden, M., Ortsäter, H., Kazenwadel, J., Calado, D.P., Ostergaard, P., Salminen, M., et al. (2018). Matrix stiffness controls lymphatic vessel formation through regulation of a GATA2-dependent transcriptional program. Nat. Commun. 9, 1511.

Geng, X., Cha, B., Mahamud, M.R., Lim, K.-C., Silasi-Mansat, R., Uddin, M.K.M., Miura, N., Xia, L., Simon, A.M., Engel, J.D., et al. (2016). Multiple mouse models of primary lymphedema exhibit distinct defects in lymphovenous valve development. Dev. Biol. 409, 218–233.

Glasgow, E., and Tomarev, S.. (1998). Restricted expression of the homeobox gene prox 1 in developing zebrafish. Mech. Dev. 76, 175–178.

Grant, C.E., Bailey, T.L., and Noble, W.S. (2011). FIMO: scanning for occurrences of a given motif. Bioinformatics 27, 1017–1018.

Grimm, L., Mason, E., Yu, H., Dudczig, S., Panara, V., Chen, T., Bower, N.I., Paterson, S., Rondon Galeano, M., Kobayashi, S., et al. (2023). Single-cell analysis of lymphatic endothelial cell fate specification and differentiation during zebrafish development. EMBO J.

Le Guen, L., Karpanen, T., Schulte, D., Harris, N.C., Koltowska, K., Roukens, G., Bower, N.I., van Impel, A., Stacker, S.A., Achen, M.G., et al. (2014). Ccbe1 regulates Vegfc-mediated induction of Vegfr3 signaling during embryonic lymphangiogenesis. Development 141, 1239–1249.

Guo, Y., Pan, W., Liu, S., Shen, Z., Xu, Y., and Hu, L. (2020). ERK/MAPK signalling pathway and tumorigenesis. Exp. Ther. Med.

Gupta, S., Stamatoyannopoulos, J.A., Bailey, T.L., and Noble, W.S. (2007). Quantifying similarity between motifs. Genome Biol. 8.

Heintzman, N.D., Stuart, R.K., Hon, G., Fu, Y., Ching, C.W., Hawkins, R.D., Barrera, L.O., Van Calcar, S., Qu, C., Ching, K.A., et al. (2007). Distinct and predictive chromatin signatures of transcriptional promoters and enhancers in the human genome. Nat. Genet. 39, 311–318.

Hobert, O. (2010). Gene Regulation: Enhancers Stepping Out of the Shadow. Curr. Biol. 20, R697–R699.

Hogan, B.M., Bos, F.L., Bussmann, J., Witte, M., Chi, N.C., Duckers, H.J., and Schulte-Merker, S. (2009). Ccbe1 is required for embryonic lymphangiogenesis and venous sprouting. Nat. Genet. 41, 396–398.

Hong, J.-W., Hendrix, D.A., and Levine, M.S. (2008). Shadow Enhancers as a Source of Evolutionary Novelty. Science (80-.). 321, 1314–1314.

Hultgren, N.W., Fang, J.S., Ziegler, M.E., Ramirez, R.N., Phan, D.T.T., Hatch, M.M.S., Welch-Reardon, K.M., Paniagua, A.E., Kim, L.S., Shon, N.N., et al. (2020). Slug regulates the Dll4-Notch-VEGFR2 axis to control endothelial cell activation and angiogenesis. Nat. Commun. 11, 5400.

van Impel, A., Zhao, Z., Hermkens, D.M.A., Roukens, M.G., Fischer, J.C., Peterson-Maduro, J., Duckers, H., Ober, E.A., Ingham, P.W., and Schulte-Merker, S. (2014). Divergence of zebrafish and mouse lymphatic cell fate specification pathways. Development 141, 1228– 1238.

Kazenwadel, J., Betterman, K.L., Chong, C.E., Stokes, P.H., Lee, Y.K., Secker, G.A., Agalarov, Y., Demir, C.S., Lawrence, D.M., Sutton, D.L., et al. (2015). GATA2 is required for lymphatic vessel valve development and maintenance. J. Clin. Invest. 125, 2879–2994.

Kazenwadel, J., Venugopal, P., Oszmiana, A., Toubia, J., Arriola-Martinez, L., Panara, V., Piltz, S.G., Brown, C., Ma, W., Schreiber, A.W., et al. (2023). A Prox1 enhancer represses haematopoiesis in the lymphatic vasculature. Nature 614, 343–348.

Koltowska, K., Lagendijk, A.K., Pichol-Thievend, C., Fischer, J.C., Francois, M., Ober, E.A., Yap, A.S., and Hogan, B.M. (2015a). Vegfc Regulates Bipotential Precursor Division and Prox1 Expression to Promote Lymphatic Identity in Zebrafish. Cell Rep. 13, 1828–1841.

Koltowska, K., Paterson, S., Bower, N.I., Baillie, G.J., Lagendijk, A.K., Astin, J.W., Chen, H., Francois, M., Crosier, P.S., Taft, R.J., et al. (2015b). Mafba Is a Downstream Transcriptional Effector of Vegfc Signaling Essential for Embryonic Lymphangiogenesis in Zebrafish. Genes Dev. 29, 1618–1630.

Küchler, A.M., Gjini, E., Peterson-Maduro, J., Cancilla, B., Wolburg, H., and Schulte-Merker, S. (2006). Development of the Zebrafish Lymphatic System Requires Vegfc Signaling. Curr. Biol. 16, 1244–1248.

Kvon, E.Z., Waymack, R., Gad, M., and Wunderlich, Z. (2021). Enhancer redundancy in development and disease. Nat. Rev. Genet. 22, 324–336.

Lam, D.D., de Souza, F.S.J., Nasif, S., Yamashita, M., López-Leal, R., Otero-Corchon, V., Meece, K., Sampath, H., Mercer, A.J., Wardlaw, S.L., et al. (2015). Partially Redundant Enhancers Cooperatively Maintain Mammalian Pomc Expression Above a Critical Functional Threshold. PLOS Genet. 11, e1004935.

Lawson, N.D., and Weinstein, B.M. (2002). In Vivo Imaging of Embryonic Vascular Development Using Transgenic Zebrafish. Dev. Biol. 248, 307–318.

Lin, F.-J., Chen, X., Qin, J., Hong, Y.-K., Tsai, M.-J., and Tsai, S.Y. (2010). Direct transcriptional regulation of neuropilin-2 by COUP-TFII modulates multiple steps in murine lymphatic vessel development. J. Clin. Invest. 120, 1694–1707.

Long, H.K., Prescott, S.L., and Wysocka, J. (2016). Ever-Changing Landscapes: Transcriptional Enhancers in Development and Evolution. Cell 167, 1170–1187.

Marino, D., Dabouras, V., Brändli, A.W., and Detmar, M. (2011). A Role for All-Trans-Retinoic Acid in the Early Steps of Lymphatic Vasculature Development. J. Vasc. Res. 48, 236–251.

Mayor, C., Brudno, M., Schwartz, J.R., Poliakov, A., Rubin, E.M., Frazer, K.A., Pachter, L.S., and Dubchak, I. (2000). Vista: Visualizing global DNA sequence alignments of arbitrary length. Bioinformatics.

Nicenboim, J., Malkinson, G., Lupo, T., Asaf, L., Sela, Y., Mayseless, O., Gibbs-Bar, L., Senderovich, N., Hashimshony, T., Shin, M., et al. (2015). Lymphatic vessels arise from specialized angioblasts within a venous niche. Nature 522, 56–61.

Niimi, K., Ueda, M., Fukumoto, M., Kohara, M., Sawano, T., Tsuchihashi, R., Shibata, S., Inagaki, S., and Furuyama, T. (2017). Transcription factor FOXO1 promotes cell migration toward exogenous ATP via controlling P2Y1 receptor expression in lymphatic endothelial cells. Biochem. Biophys. Res. Commun. 489, 413–419.

Niimi, K., Kohara, M., Sedoh, E., Fukumoto, M., Shibata, S., Sawano, T., Tashiro, F., Miyazaki, S., Kubota, Y., Miyazaki, J., et al. (2019). FOXO1 regulates developmental lymphangiogenesis by upregulating CXCR4 in the mouse-tail dermis. Development.

Okuda, K.S., Astin, J.W., Misa, J.P., Flores, M. V., Crosier, K.E., and Crosier, P.S. (2012). Lyve1 expression reveals novel lymphatic vessels and new mechanisms for lymphatic vessel development in zebrafish. Dev.

Osterwalder, M., Barozzi, I., Tissières, V., Fukuda-Yuzawa, Y., Mannion, B.J., Afzal, S.Y., Lee, E.A., Zhu, Y., Plajzer-Frick, I., Pickle, C.S., et al. (2018). Enhancer redundancy provides phenotypic robustness in mammalian development. Nature 554, 239–243.

Park, D.-Y., Lee, J., Park, I., Choi, D., Lee, S., Song, S., Hwang, Y., Hong, K.Y., Nakaoka, Y., Makinen, T., et al. (2014). Lymphatic regulator PROX1 determines Schlemm’s canal integrity and identity. J. Clin. Invest. 124, 3960–3974.

Pham, V.N., Lawson, N.D., Mugford, J.W., Dye, L., Castranova, D., Lo, B., and Weinstein, B.M. (2007). Combinatorial function of ETS transcription factors in the developing vasculature. Dev. Biol. 303, 772–783.

Pistocchi, A., Gaudenzi, G., Carra, S., Bresciani, E., Del Giacco, L., and Cotelli, F. (2008). Crucial role of zebrafish prox1in hypothalamic catecholaminergic neurons development. BMC Dev. Biol. 8, 27.

Quillien, A., Abdalla, M., Yu, J., Ou, J., Zhu, L.J., and Lawson, N.D. (2017). Robust Identification of Developmentally Active Endothelial Enhancers in Zebrafish Using FANS-Assisted ATAC-Seq. Cell Rep.

Scallan, J.P., Knauer, L.A., Hou, H., Castorena-Gonzalez, J.A., Davis, M.J., and Yang, Y. (2021). Foxo1 deletion promotes the growth of new lymphatic valves. J. Clin. Invest. 131.

Shin, M., Male, I., Beane, T.J., Villefranc, J.A., Kok, F.O., Zhu, L.J., and Lawson, N.D. (2016). Vegfc acts through ERK to induce sprouting and differentiation of trunk lymphatic progenitors. Development 143, 3785–3795.

Shin, M., Nozaki, T., Idrizi, F., Isogai, S., Ogasawara, K., Ishida, K., Yuge, S., Roscoe, B., Wolfe, S.A., Fukuhara, S., et al. (2019). Valves Are a Conserved Feature of the Zebrafish Lymphatic System. Dev. Cell 51, 374–386.e5.

Spitz, F., and Furlong, E.E.M. (2012). Transcription factors: from enhancer binding to developmental control. Nat. Rev. Genet. 13, 613–626.

Srinivasan, R.S., Geng, X., Yang, Y., Wang, Y., Mukatira, S., Studer, M., Porto, M.P.R., Lagutin, O., and Oliver, G. (2010). The nuclear hormone receptor Coup-TFII is required for the initiation and early maintenance of Prox1 expression in lymphatic endothelial cells. Genes Dev. 24, 696–707.

Tai-Nagara, I., Hasumi, Y., Kusumoto, D., Hasumi, H., Okabe, K., Ando, T., Matsuzaki, F., Itoh, F., Saya, H., Liu, C., et al. (2020). Blood and lymphatic systems are segregated by the FLCN tumor suppressor. Nat. Commun. 11, 6314.

Tang, T., Shi, Y., Opalenik, S.R., Brantley-Sieders, D.M., Chen, J., Davidson, J.M., and Brandt, S.J. (2006). Expression of the TAL1/SCL transcription factor in physiological and pathological vascular processes. J. Pathol. 210, 121–129.

Thomas, P.D., Kejariwal, A., Guo, N., Mi, H., Campbell, M.J., Muruganujan, A., and Lazareva-Ulitsky, B. (2006). Applications for protein sequence-function evolution data: mRNA/protein expression analysis and coding SNP scoring tools. Nucleic Acids Res. 34, W645–W650.

De Val, S., Chi, N.C., Meadows, S.M., Minovitsky, S., Anderson, J.P., Harris, I.S., Ehlers, M.L., Agarwal, P., Visel, A., Xu, S.-M., et al. (2008). Combinatorial Regulation of Endothelial Gene Expression by Ets and Forkhead Transcription Factors. Cell 135, 1053–1064.

Varshney, G.K., Carrington, B., Pei, W., Bishop, K., Chen, Z., Fan, C., Xu, L., Jones, M., LaFave, M.C., Ledin, J., et al. (2016). A high-throughput functional genomics workflow based on CRISPR/Cas9-mediated targeted mutagenesis in zebrafish. Nat. Protoc. 11, 2357–2375.

Veldman, M.B., and Lin, S. (2012). Etsrp/Etv2 Is Directly Regulated by Foxc1a/b in the Zebrafish Angioblast. Circ. Res. 110, 220–229.

Villefranc, J.A., Amigo, J., and Lawson, N.D. (2007). Gateway compatible vectors for analysis of gene function in the zebrafish. Dev. Dyn. 236, 3077–3087.

Wigle, J.T., and Oliver, G. (1999). Prox1 function is required for the development of the murine lymphatic system. Cell.

Wong, E.S., Zheng, D., Tan, S.Z., Bower, N.I., Garside, V., Vanwalleghem, G., Gaiti, F., Scott, E., Hogan, B.M., Kikuchi, K., et al. (2020). Deep conservation of the enhancer regulatory code in animals. Science (80-.). 370, eaax8137.

Yaniv, K., Isogai, S., Castranova, D., Dye, L., Hitomi, J., and Weinstein, B.M. (2006). Live imaging of lymphatic development in the zebrafish. Nat. Med. 12, 711–716.

Yoshimatsu, Y., Yamazaki, T., Mihira, H., Itoh, T., Suehiro, J., Yuki, K., Harada, K., Morikawa, M., Iwata, C., Minami, T., et al. (2011). Ets family members induce lymphangiogenesis through physical and functional interaction with Prox1. J. Cell Sci. 124, 2753–2762.

