## Supplementary figures and images for "Multiple *cis*-regulatory elements control *prox1a* expression in distinct lymphatic vascular beds"

### Supplemental Figure 1

#### Supplementary Figure 1

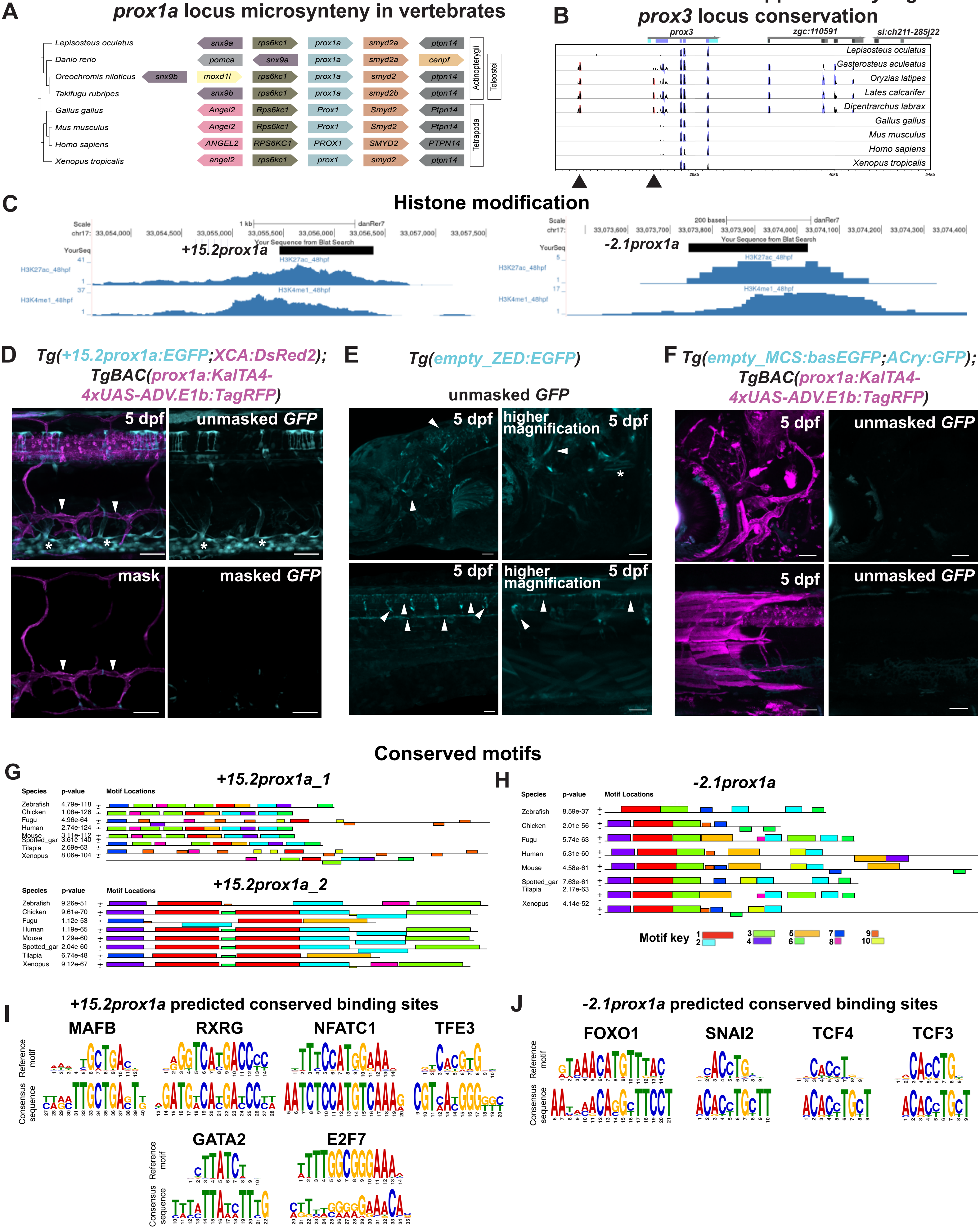

### Supplemental Figure 2

Supplementary Figure 2

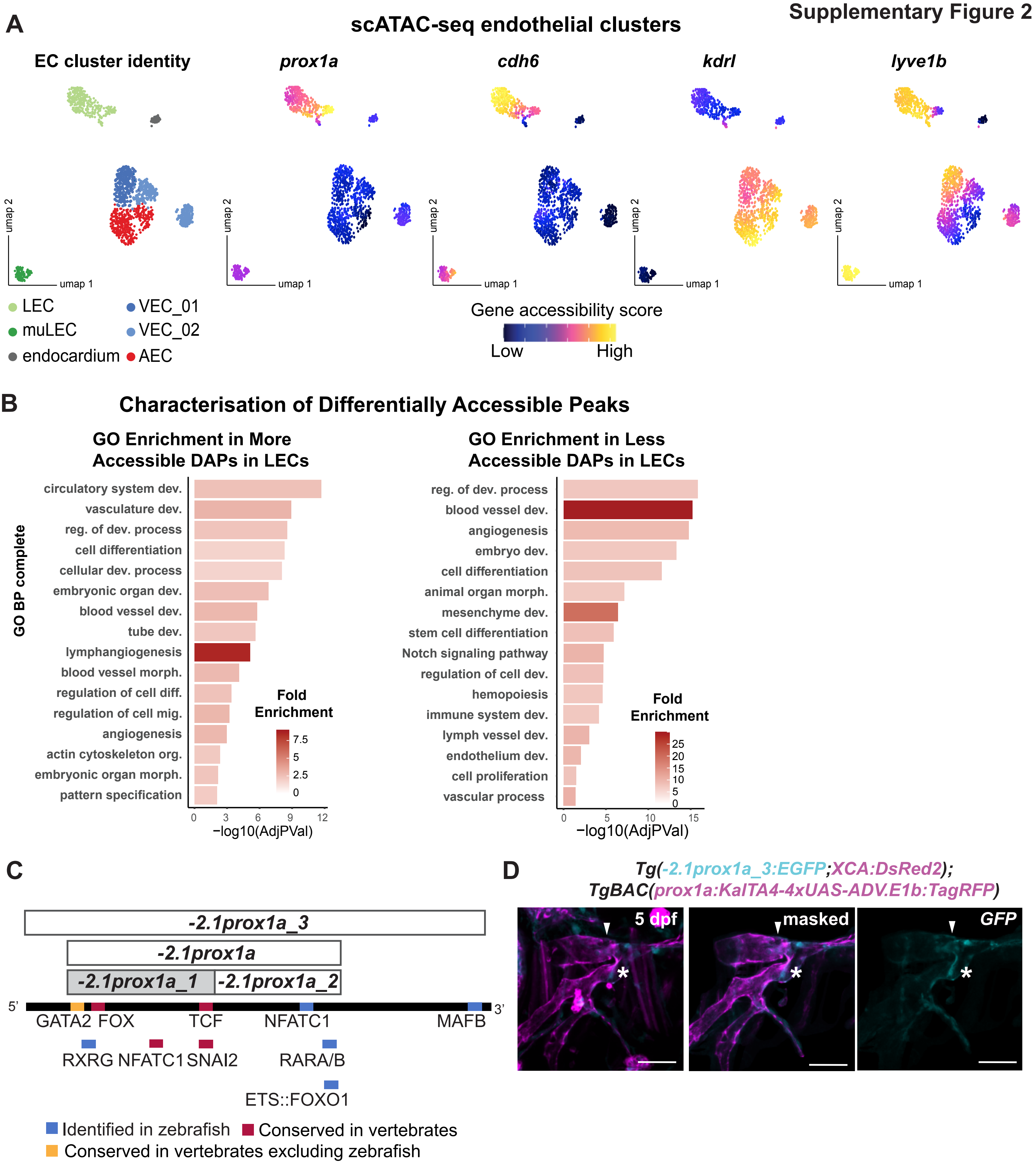

### Supplemental Figure 3

#### Supplementary Figure 3

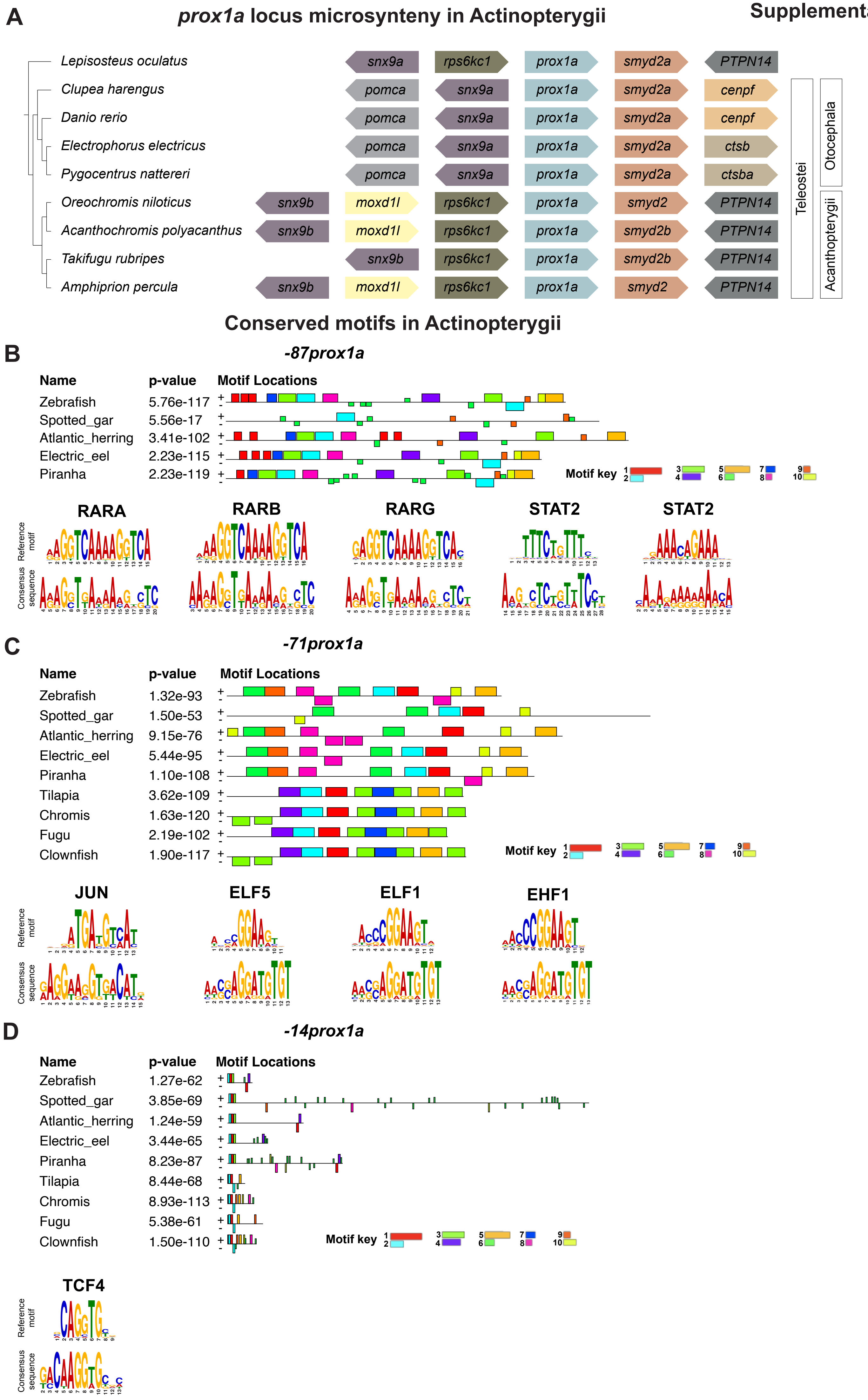

### Supplemental Figure 4

#### Supplementary Figure 4

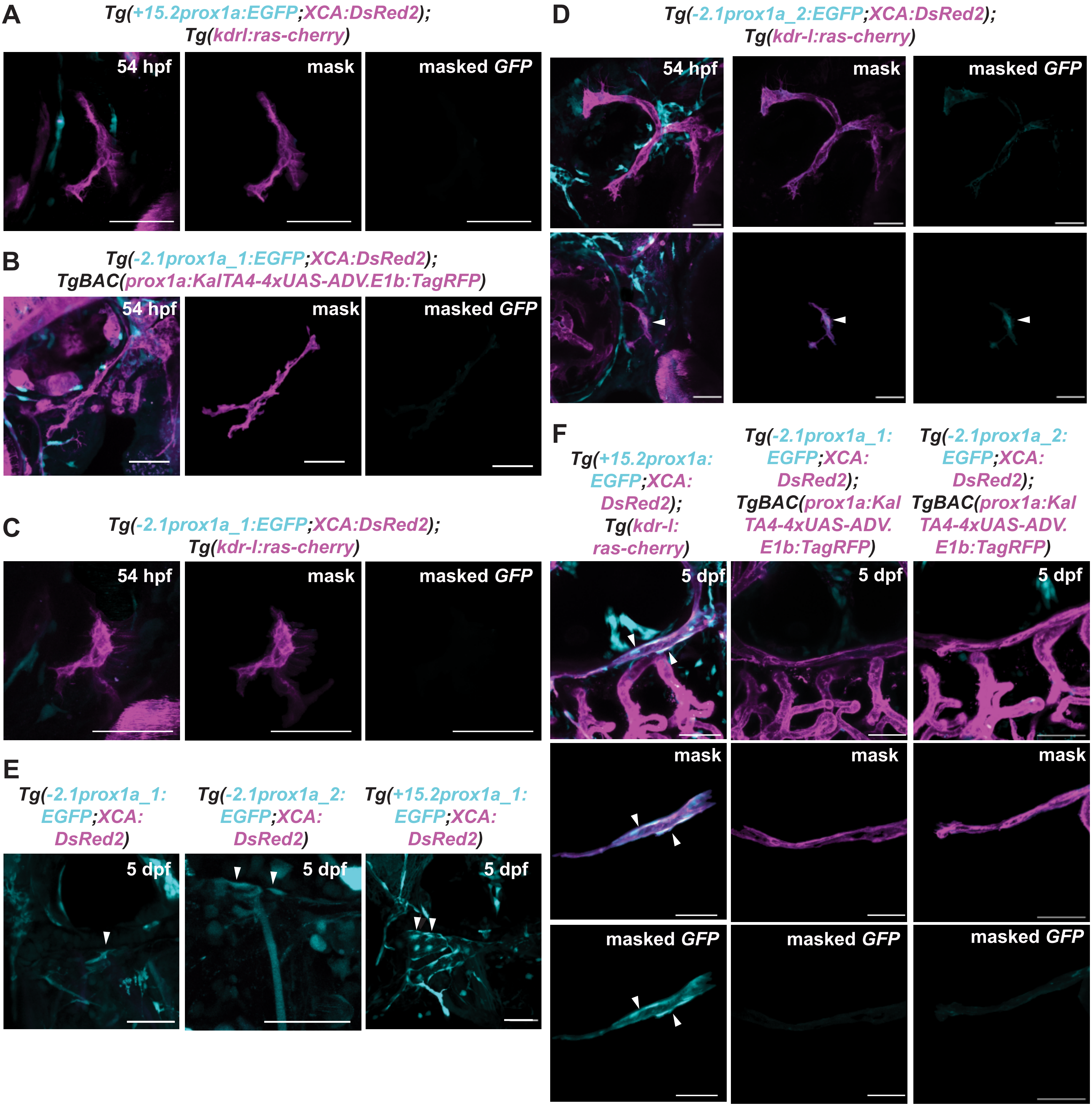

### Supplemental Figure 5

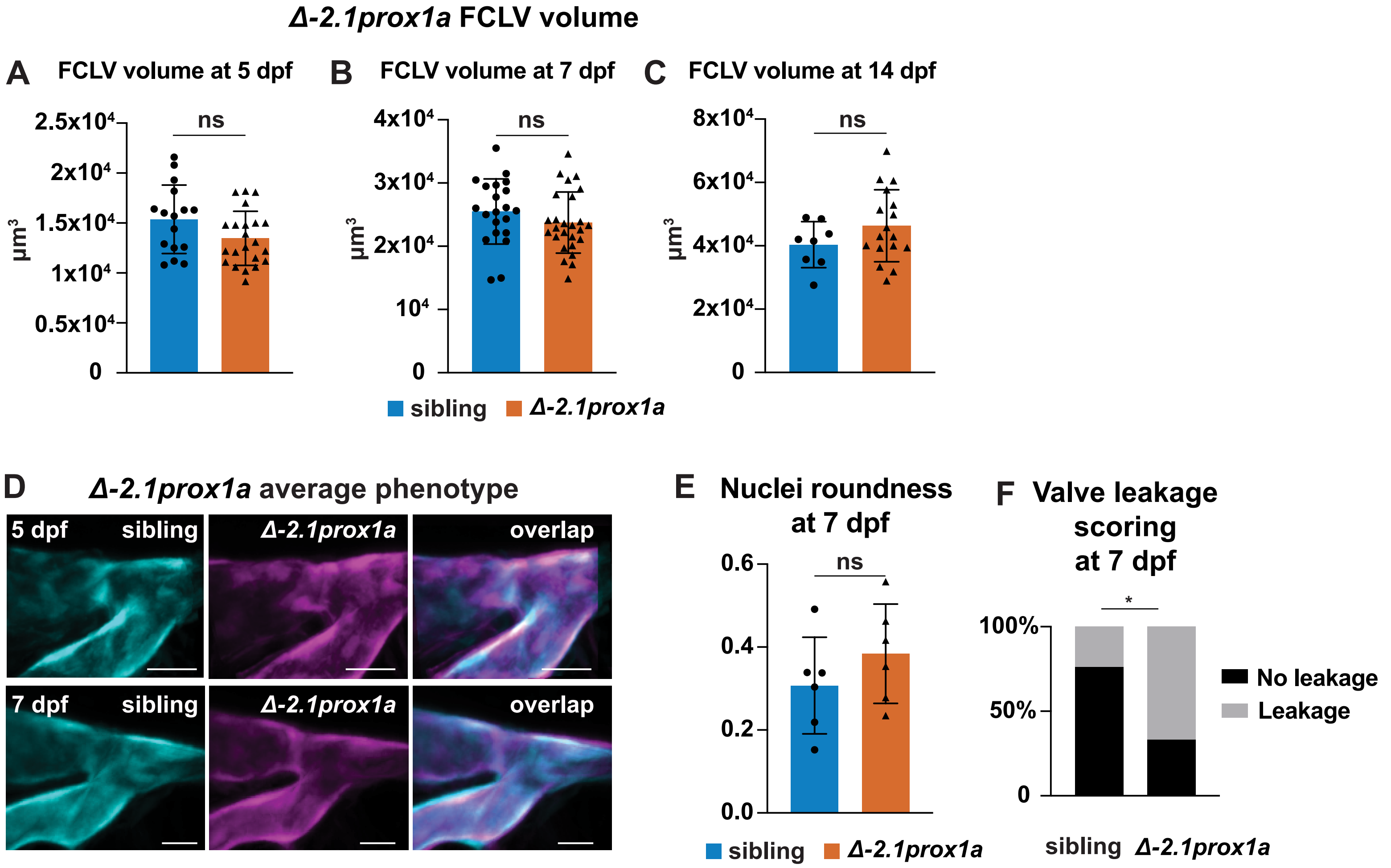
